# FIGNL1 inhibits homologous recombination in BRCA2 deficient cells by dissociating RAD51 filaments

**DOI:** 10.1101/2024.11.03.621741

**Authors:** Raviprasad Kuthethur, Ananya Acharya, Satheesh Kumar Sengodan, Carmen Fonseca, Nupur Nagar, Safa Nasrin VZ, Oluwakemi Ibini, Eleni-Maria Manolika, Kelly de Koning, Stefan Braunshier, Julien Dessapt, Amélie Fradet-Turcotte, Joyce H.G. Lebbink, Roland Kanaar, Krishna Mohan Poluri, Shyam K. Sharan, Petr Cejka, Arnab Ray Chaudhuri

## Abstract

Homologous recombination (HR) deficiency upon BRCA2 loss arises from defects in the formation of RAD51 nucleoprotein filaments. Here, we demonstrate that loss of the anti-recombinase FIGNL1 retains RAD51 loading at DNA double-stranded breaks (DSBs) in BRCA2-deficient cells, leading to genome stability, HR proficiency, and viability of BRCA2-deficient mouse embryonic stem cells. Mechanistically, we directly show that strand invasion and subsequent HR defects upon BRCA2 loss primarily arises from the unrestricted removal of RAD51 from DSB sites by FIGNL1, rather than from defective RAD51 loading. Furthermore, we identify that the MMS22L-TONSL complex interacts with FIGNL1 and is critical for HR in BRCA2/FIGNL1 double-deficient cells. These findings identify a pathway for tightly regulating RAD51 activity to promote efficient HR, offering insights into mechanisms of chemoresistance in BRCA2-deficient tumors.

## INTRODUCTION

The homologous recombination (HR) pathway is essential for accurate repair of double-stranded breaks (DSBs) to prevent mutagenesis and genome instability (*1*). The tumor suppressor protein BRCA2 is a critical factor required for HR process and directly interacts with the central recombinase RAD51. This interaction in turn mediates the nucleation and stabilization of RAD51 nucleoprotein filaments on end-resected single-stranded DNA (ssDNA) by displacing RPA (*2–4*). This step is crucial for downstream DNA strand invasion and the synthesis steps of the HR process. Consistent with this vital function of BRCA2, its loss results in defects in RAD51 accumulation at DSBs, thus disrupting HR and leading to subsequent genome instability and loss of cellular survival (*5–11*). Tumor cells with BRCA2 mutations also exhibit defects in RAD51 loading and HR deficiencies (*12–15*) and serve as potential targets for chemotherapeutics that induce DSBs, such as poly(adenosine-diphosphate-ribose) polymerase inhibitors (PARPi) and DNA crosslinking agents (*15–17*). Although these treatment strategies for BRCA2-deficient tumors have yielded significant success, a subset of these tumors acquire resistance to these therapies through compensatory mechanisms, including reversion mutations and replication fork stability (*13, 18, 19*). However, no molecular mechanisms of restoration of RAD51 loading and HR have been identified upon BRCA2 loss, underscoring its essential role in mediating HR.

In addition to BRCA2, the MMS22L-TONSL heterodimeric complex regulates HR by controlling RAD51 dynamics at replication-induced DSBs (*20–22*). This complex interacts with RPA and promote RAD51-mediated strand invasion, thereby stimulating HR (*22*). MMS22L contains RAD51 interaction motifs that facilitate RAD51 loading upon DNA damage (*22*). In addition to interacting with MMS22L, TONSL also interacts with ASF1 and CAF1 and recognizes H4K20me0 at DSBs, resulting in MMS22L-mediated loading of RAD51 at DSBs (*21, 23, 24*). Loss of MMS22L-TONSL, however, does not affect the localization of BRCA2 to DSBs, suggesting that it may function either downstream of BRCA2 or in a separate pathway for controlling RAD51-mediated HR (*21*). Nevertheless, the coordinated action of these two pathways to ensure efficient HR remains poorly understood.

The loading and stabilization of RAD51 necessitates stringent regulation, as excessive accumulation of RAD51 on chromatin is associated with genome instability and cell death (*25, 26*). This regulation is achieved through the dissociation of RAD51 from DNA, mediated by multiple helicases and translocases, including the RECQ family helicases, PARI, RTEL1, and FBH1 (*27–32*). These translocases and helicases disassemble RAD51 at various stages of the HR process, thereby ensuring its efficiency. FIGNL1 (fidgetin-like 1), a member of the large AAA+ ATPase family, plays a significant role in the maintenance of genome stability (*33*). Depletion of FIGNL1 results in excessive accumulation of RAD51, leading to genome instability manifested as persistent ultrafine chromosome bridges (*33*). FIGNL1 and its interaction with FIRRM/FLIP are essential for DNA cross-link repair and HR by facilitating the dissociation of RAD51 (*34–39*). FIGNL1 is also crucial for meiotic recombination, as its absence results in impaired spermatogenesis arising from hyperaccumulation of RAD51 (*40, 41*). This recent evidence underscores the importance of FIGNL1 in the regulation of RAD51 dynamics during HR. However, the molecular mechanisms by which the intricate balance of RAD51 loading and dissociation is regulated, and its implications for HR fidelity and genome stability, remain unresolved.

## RESULTS

### FIGNL1 loss restores RAD51 loading at DSBs upon BRCA2 deficiency

Loss of FIGNL1 increases chromatin association of RAD51 and is not compatible with cellular viability (*40, 42*). Therefore, we genetically inactivated FIGNL1 using CRISPR-CAS9 in RPE1 hTERT p53^-/-^ cells harboring a doxycycline (Dox) inducible shRNA against BRCA2 (WT or shBRCA2) (*43*) (**Fig.S1A-B**). FIGNL1 inactivation was also performed in *Brca2*^−/−^*; Trp53*^−/−^ mouse mammary tumor cell line KB2P1.21 (*Brca2*^−/−^) and *Brca2* reconstituted line (WT), derived from *K14cre;Trp53*^F/F^*;Brca2*^F/F^ mouse model for *BRCA2*-mutated breast cancer (*44*) (**Fig. S1C-D**). Inactivation of FIGNL1 in a p53-deficient background yielded viable cells, and cell cycle analysis of these lines revealed no significant changes in cell cycle profiles, suggesting that genome instability arising from loss of FIGNL1 is well tolerated upon p53 deficiency (**Fig.S1E-F**). We then tested for the effects of RAD51 accumulation in the presence or absence of DSBs upon loss of FIGNL1 in BRCA2-proficient or BRCA2-deficient conditions. To this end, we treated cells with ionizing radiation (IR) to induce DSBs and probed for chromatin-bound RAD51 accumulation into foci using quantitative immunofluorescence-based cytometry (QIBC) (*45*). As expected, loss of FIGNL1 in both RPE1 and mouse mammary tumor lines resulted in higher levels of RAD51 accumulation on chromatin. The increase of chromatin bound RAD51 was apparent under untreated conditions and was significantly enhanced exposure to IR (**Fig.1A and Fig.S2A**). Loss of BRCA2 resulted in a significantly defective accumulation of RAD51 in foci along expected lines (**Fig.1A and Fig.S2A**). Surprisingly, loss of FIGNL1 in BRCA2-deficient cells significantly rescued the RAD51 focus formation defect observed in BRCA2-deficient cells alone (**Fig.1A and Fig.S2A**) upon the induction of DSBs. Since IR treatment can result in multiple types of DNA damage, including DSBs, we next tested whether the loss of FIGNL1 results in higher RAD51 focus formation upon induction of site-specific DSBs. To test this, we stably expressed restriction enzyme AsiSI in mouse mammary tumor lines, in which site-specific DSBs across the genome can be induced upon co-treatment with Dox and 4-hydroxytamoxifen (4-OHT) (**Fig.1B and Fig.S2B)** (*46*). Induction of AsiSI resulted in similar levels of DSBs in all cell lines, measured by analyzing 53BP1 foci using QIBC (**Fig.S2B)**. In line with our data on IR-induced DSBs, induction of site-specific DSBs in mouse mammary tumor lines also exhibited significantly higher levels of RAD51 foci formation upon loss of FIGNL1 when compared to WT cells. Loss of BRCA2 abrogated RAD51 foci formation upon induction of site-specific breaks, which was rescued back to WT levels upon concomitant loss of FIGNL1 in these cells, in line with IR-induced DSBs (**Fig.1B).** Furthermore, we tested whether this effect was cell line- and organism-specific by downregulating either BRCA2, FIGNL1, or both in U2OS-DIvA cells treated with 4-OHT to induce site-specific DSBs by AsiSI (*47*). Consistent with our data from mouse mammary tumors, loss of FIGNL1 in BRCA2 deficient U2OS cells significantly rescued the RAD51 foci formation defects observed upon BRCA2 loss (**Fig.S2C)**. Taken together, these data suggest that the RAD51 accumulation defects at DSBs observed upon the loss of BRCA2 could at least in part result from RAD51 removal by FIGNL1, and not defective RAD51 loading.

**Figure 1:**
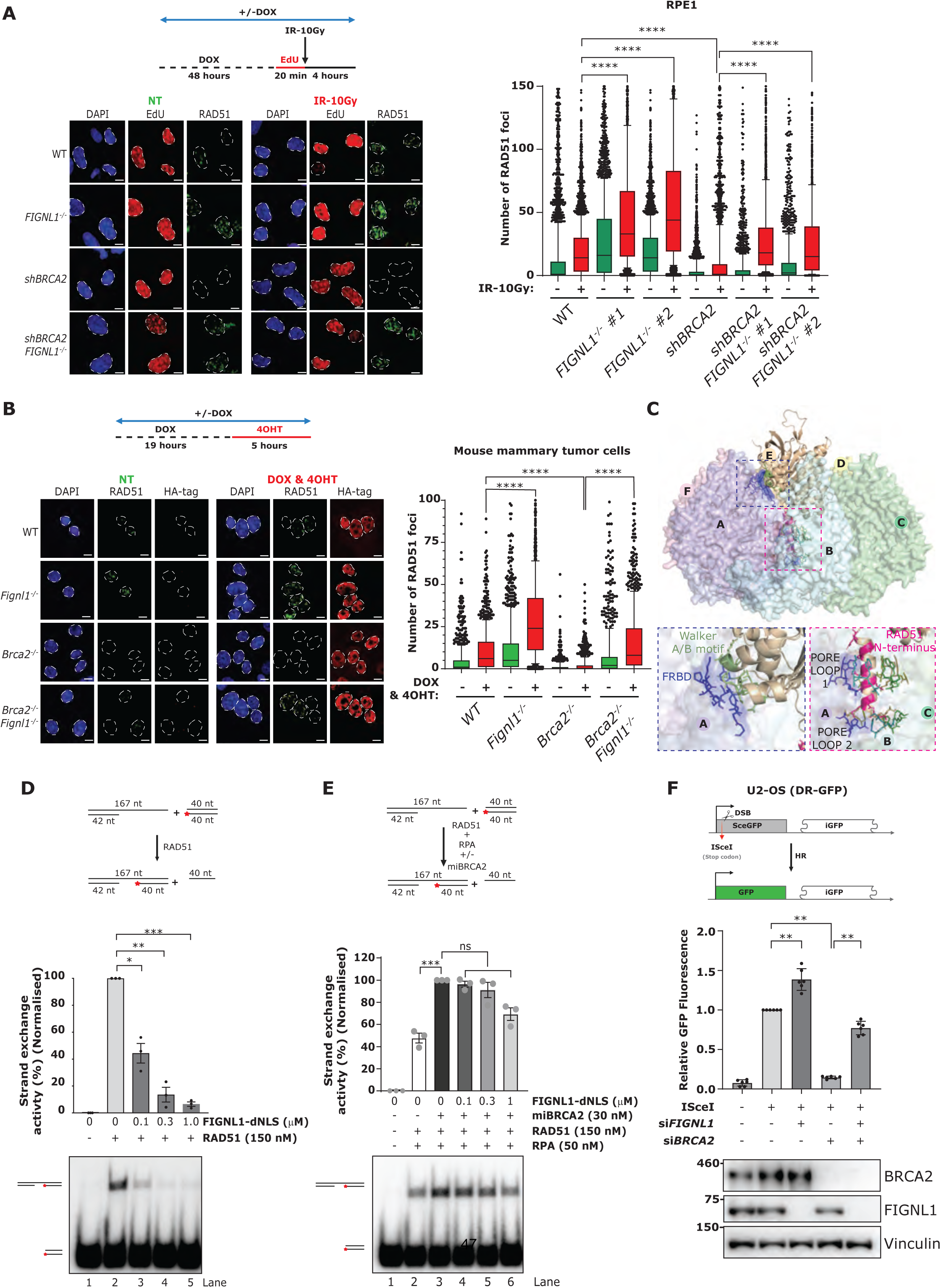
Loss of FIGNL1 in BRCA2 deficient cells rescues RAD51 mediated HR. (A) Top: Schematics showing the experimental setup for the high-content, automated RAD51 immunofluorescence (IF) in human RPE1 cells. 2µg/ml of doxycycline for 48 hours to induce the knock-down of BRCA2, followed by EdU labelling and induction of DSB via 10Gy of X-ray treatment. Left: Representative IF images showing EdU and RAD51 foci in untreated (NT) and ionizing radiation treated (IR) cells; scale bar: 20µm. Right: Automated Quantitative Image Based Cytometry (QIBC) plot of the RAD51 foci per nucleus; scatter box plot showing the foci distribution between 10-90th percentile from 2000 cells (experiment was repeated 3 times). P-values from Mann-Whitney test (two-tailed) comparing between indicated groups (****: <0.0001). (B) Top: Schematics showing the experimental setup for RAD51-IF in mouse mammary tumor cells expressing AsiSI with HA-tag. 24 hours of doxycycline treatment to induce AsiSI-HA expression with last 5 hours to translocate it to the nucleus by addition of 4-hydroxytamoxifen (4-OHT). Left: Representative IF images of the RAD51 and HA-tag in un-induced (NT) and induced (DOX &4 OHT) cells; scale bar: 20µm. Right: Automated QIBC plots of the RAD51 foci per nucleus; scatter box plot showing the foci distribution between 10-90th percentile from 1500 cells (experiment was repeated 3 times). P-values from Mann-Whitney test (two-tailed) comparing between indicated groups (****: <0.0001). (C) Top: The side and zoom-in view of the predicted FIGNL1-RAD51 complex. The FIGNL1 monomers are shown as surface representation (Chain A: Light blue; Chain B: Pale cyan; Chain C: Pale green; Chain D: Light yellow; Chain E: Light Orange; Chain F: Light pink) while RAD51 (beige) is shown in cartoon representation. Bottom: The zoom-in views represent the interacting regions of FIGNL1 and RAD51. Blue inset: the FIGNL1 FRBD (dark blue) interacts with a loop connecting the RAD51 Walker A/B motifs (green). Pink inset: two Pore Loops present in the FIGNL1 ATPase region (dark blue) interact with the RAD51 N-terminal domain (pink). (D) Top: Schematics of the DNA strand exchange assays. The assays were performed by adding 150nM purified RAD51 to the DNA substrates in the presence of increasing concentrations of purified FIGNL1-dNLS. The red asterisk indicates the position of the radioactive label. Bottom: Percentage normalized quantifications showing the band intensity of the exchanged DNA; error bars are indicative of mean±SEM from 3 independent experiments (n=3) P-values from Dunnett’s multiple comparison One-Way ANOVA comparing between indicated groups (*: <0.05; **: <0.01; ***: <0.001). (E) Top: Schematics of the DNA strand exchange assays. The assays were performed by adding 150nM purified RAD51 and 50nM RPA to the DNA substrates. The reactions were then supplemented with 30nM miBRCA2 and increasing concentrations of purified FIGNL1-dNLS. The red asterisk indicates the position of the radioactive label. Bottom: Percentage normalized quantifications showing the band intensity of the exchanged DNA; error bars are indicative of mean±SEM from 3 independent experiments (n=3). P-values from Dunnett’s multiple comparison One-Way ANOVA comparing between indicated groups (***: <0.001). (F) Top: Schematics of DR-GFP assay showing the strategy behind the experiment. Mutant SceGFP containing ISceI recognition site and stop codon is repaired by the downstream iGFP to restore the GFP fluorescence in the cells. Center: U2-OS cells with a stable integrated DR-GFP system were transfected for 72 hours with the indicated siRNA’s and underwent 48 hours of ISceI treatment to measure the levels of GFP fluorescence relative to the WT cells undergoing ISceI treatment. Error bars are representative of the mean±SD from 3 independent experiments (n=3). P-values from unpaired t-test comparing between indicated groups (**: >0.01). Bottom: Western blot analysis showing the levels of FIGNL1 and BRCA2 from the corresponding cells used for FACS analysis.

FIGNL1 interacts with FIRRM/FLIP and can disassemble RAD51 nucleoprotein filaments (*34, 35, 39*). To directly test whether the DNA binding of FIGNL1 is required for this function, we purified an N-terminal truncated fragment of FIGNL1 (FIGNL1-dNLS) lacking 1-120 residues which disrupts the binding with its interaction partner FIRRM/FLIP but is biochemically functional (**Fig.S3A**) (*35*). Our electrophoretic mobility shift assays revealed that FIGNL1-dNLS binds both ssDNA and double-stranded DNA (dsDNA) with weak affinity at lower concentrations, with DNA binding observed only at high concentrations of FIGNL1 (**Fig.S3B**). Furthermore, the addition of the ssDNA binding protein RPA did not enhance the binding efficiency of FIGNL1 to ssDNA (**Fig.S3C-D**) suggesting that FIGNL1 probably disrupts RAD51 nucleofilaments through direct interaction with the RAD51 protein. FIGNL1 has previously been shown to interact with RAD51 (*38*). Our *in vitro* interaction assay with purified FIGNL1-dNLS and RAD51 and pulldown assays with reconstituted FIGNL1 from RPE1 cells indeed revealed a strong interaction between FIGNL1 and RAD51 (**Fig.S3E-G**). Next, to further gain further insights into the molecular interactions involving RAD51 binding to FIGNL1, we used AlphaFold 3 to predict the structure of FIGNL1, which was previously shown to exist as a hexamer in solution (*35, 42*). Indeed, AlphaFold 3 predictions showed a hexameric structure for FIGNL1 involving interactions between the N-terminal NLS, ATPase, and C-terminal V domains (**Fig.S3H-S3I, Table.1A**). The monomer-monomer interaction mainly comprises of two binding pockets. The first pocket is the NLS-V domain pocket, in which the NLS domain of one monomer interacts with the V domain of the neighboring monomer. The second pocket involves interaction between the ATPase domains of all six FIGNL1 monomers, thus forming the hexameric pore region (**Fig.S3I**). Next, we generated an AlphaFold3 model for the RAD51 interaction with the FIGNL1 hexamer. The predicted structure indicated that RAD51 can be embedded into the hexameric pores of FIGNL1 via its N-terminal end **(Fig.1C, Table.1B)**. Assessment of the molecular architecture inferred three regions/motifs of FIGNL1-a) the <u>F</u>IGNL1-<u>R</u>AD51-<u>B</u>inding-<u>D</u>omain (FRBD) region, b) Pore Loop 1, and c) Pore Loop 2 of the ATPase domain. The FRBD region of FIGNL1 was predicted to interact with the two loops present between the Walker A (residue 127-134) and Walker B (residues 218–222) motifs of RAD51 **(Fig.1C, bottom left, Table.1B).** The Pore loops 1 and 2 were predicted to interact with the loops present between Walker A/Walker B motifis and the N-terminal helix of RAD51, thus contributing to the formation of the FIGNL1-RAD51 complex **(Fig.1C, Table.1B)**. Furthermore, we observed that all six monomeric units of FIGNL1 interacted with RAD51 **(Fig.1C, bottom right panel, Table.1B)**. Our model suggests that FIGNL1 interacts with RAD51 through multiple domains, including FRBD and ATPase domains.

**Table 1A:** Intermolecular interactions between FIGNL1 monomers.

| FIGNL1 hexamer | Interacting domains |
| --- | --- |
| Interacting region 1 | NLS domain [2-14, 40-50] – V domain [606-636; 646-656; 666-671] |
| Interacting region 2 | ATPase domain [461-481; 500-534; 546-562] – ATPase domain [461-481; 500-534; 546-562] |

**Table 1B:** Intra-molecular interactions between FIGNL1 and RAD51.

| FIGNL1-RAD51 | Interacting domains |
| --- | --- |
| Interacting region 1 | FRBD region [295-310; 315-320; 334-345]; (FIGNL1) – Loops between Walker motifs [146-150; 176-184] (RAD51) |
| Interacting region 2 | (Pore loop 1) [472-475] (FIGNL1) – N-terminal α-helix [1-10] (RAD51) |
| Interacting region 3 | (Pore loop 2) [513-517] (FIGNL1) – N-terminal α-helix [1-10] (RAD51) |

**Table 1C:**
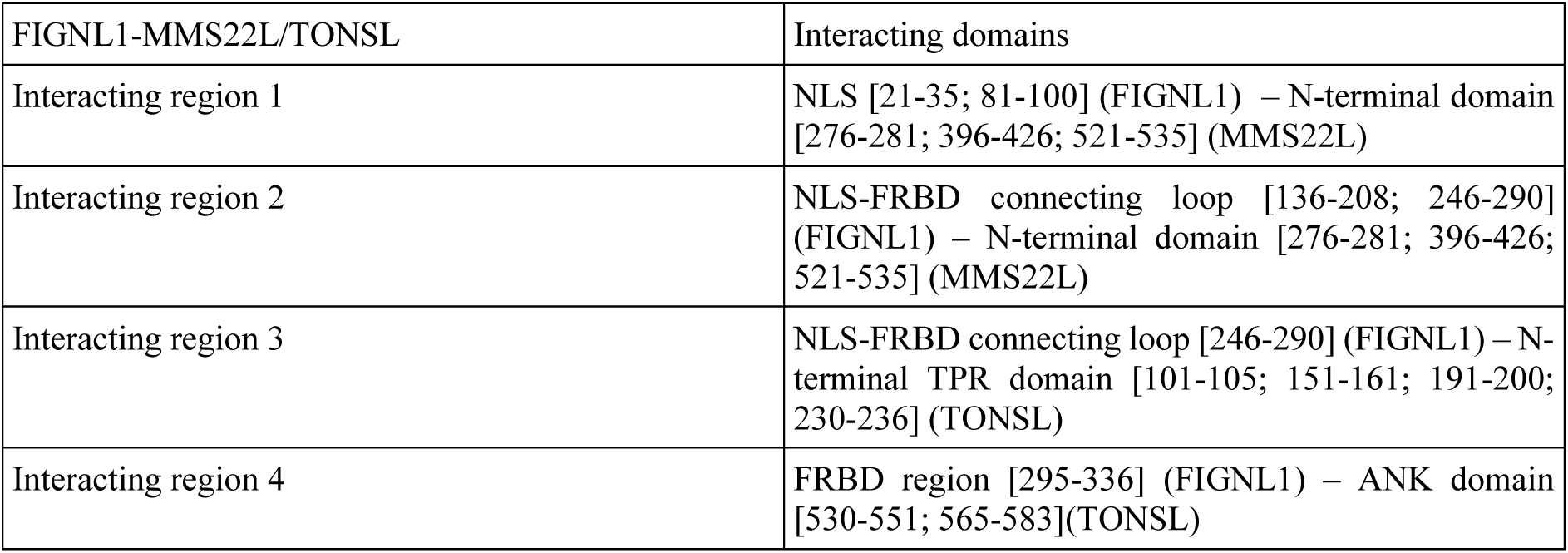
Intra-molecular interactions between FIGNL1 and MMS22L-TONSL.

| FIGNL1-MMS22L/TONSL | Interacting domains |
| --- | --- |
| Interacting region 1 | NLS [21-35; 81-100] (FIGNL1) – N-terminal domain [276-281; 396-426; 521-535] (MMS22L) |
| Interacting region 2 | NLS-FRBD connecting loop [136-208; 246-290] (FIGNL1) – N-terminal domain [276-281; 396-426; 521-535] (MMS22L) |
| Interacting region 3 | NLS-FRBD connecting loop [246-290] (FIGNL1) – N-terminal TPR domain [101-105; 151-161; 191-200; 230-236] (TONSL) |
| Interacting region 4 | FRBD region [295-336] (FIGNL1) – ANK domain [530-551; 565-583](TONSL) |

Next, to assess whether FIGNL1 disassembles RAD51 from ssDNA, we performed nuclease protection assays. Without RAD51, the addition of the DNA2 nuclease resulted in a substantial DNA degradation. When RAD51 was included, DNA was largely protected, indicative of RAD51 binding to DNA, which prevents the DNA2 nuclease from degrading it. Our data showed that the addition of FIGNL1-dNLS to RAD51 bound ssDNA resulted in decreased protection of ssDNA in a concentration-dependent manner (**Fig.S3J**), indicative of RAD51 dissociation from ssDNA in the presence of FIGNL1-dNLS. and from DNA degradation by DNA2 was tested. We then tested whether the dissociation of RAD51 from ssDNA by FIGNL-dNLS also affected its DNA strand exchange activity. As shown previously (*5*), RAD51 catalyzes the exchange of a 3’-tailed oligonucleotide-based substrate with a radiolabeled dsDNA donor (**Fig.1D)**. Addition of increasing concentrations of FIGNL1-dNLS significantly inhibited the formation of the strand exchange product, suggesting that the disruption of RAD51 filaments by FIGNL1 results in DNA strand exchange defects (**Fig.1D)**. Our immunofluorescence data on RAD51 loading at DSBs suggest that the presence of BRCA2 could potentially prevent FIGNL1 mediated RAD51 dissociation (**Fig.1A-B and S2A-C)**. Therefore, we directly tested this hypothesis *in vitro* by biochemically reconstituting the strand exchange reaction in the presence of RPA and mini BRCA2 (miBRCA2), a truncated form of BRCA2 containing the BRC1 and BRC2 motifs, DNA-binding, and C-terminal domains of BRCA2 (*48*) (**Fig.S3K**). As expected, addition of miBRCA2 stimulated the strand exchange reaction in the presence of RAD51 and RPA (**Fig.1E**). Interestingly, the presence of BRCA2 in the reaction significantly prevented the inhibition of DNA strand exchange by FIGNL1-dNLS compared to RAD51 alone (**Fig.1E and D**). Taken together, these data suggest that BRCA2 efficiently counteracts FIGNL1 mediated RAD51 dissociation to allow efficient HR. Finally, we tested whether loss of FIGNL1 indeed results in the restoration of functional HR in BRCA2 deficient cells. We performed siRNA-mediated downregulation of BRCA2, FIGNL1, and BRCA2/FIGNL1 in U2OS DR-GFP cells (*49*). Expression of ISceI endonuclease in FIGNL1 deficient cells resulted in a slight but significant increase in HR efficiency (**Fig.1F)**. Downregulation of BRCA2 resulted in a significant decrease in HR efficiency as expected. Consistent with our data on RAD51 loading, FIGNL1 deficiency significantly rescued HR defects observed in BRCA2 deficient cells. Taken together, our data strongly suggest that the HR defects upon BRCA2 loss result from the dissociation of RAD51 from DNA by FIGNL1 rather than from loading and stabilization defects of RAD51 nucleofilaments.

### Loss of FIGNL1 rescues lethality of BRCA2 deficient ESC and resistance to chemotherapeutic drugs

Genetic knockouts of either *Brca2* or *Fignl1* in mouse embryonic stem cells (mESCs) are incompatible with cellular survival possibly because of defects in the tight control of HR (*9, 40, 42, 50*). As our data suggest that loss of FIGNL1 in BRCA2 deficient cells rescues HR, we tested whether FIGNL1 deficiency can rescue the lethality of BRCA2 deficient mESCs. We generated FIGNL1 deficient PL2F7 mESCs with one null and one conditional allele of *Brca2* (*Brca2^f/−^*) (*51*) by targeting short hairpin RNAs (shRNAs) and selecting colonies that showed residual levels of FIGNL1 (**Fig. 2A and S4A**). Upon transfection of *Brca2^f/−^*mESCs with CRE and selection in HAT medium, very few resistant colonies were obtained. However, 100% of these colonies were found to be *Brca2^f/−^* rather than *Brca2*-null, indicating an essential role of BRCA2 in ESC viability (**Fig. 2B**). Strikingly, targeting FIGNL1 with two separate shRNAs yielded 37.1% and 53.2% HAT-resistant *Brca2*-null colonies after CRE expression (**Fig. 2B and S4B**). Next, we tested whether *Brca2/Fignl1* double-deficient mESCs were proficient in RAD51 loading upon the induction of DSBs. Consistent with our data from mouse and human cell lines, irradiation-induced RAD51 focus formation was significantly rescued in *Brca2/Fignl1* double-deficient mESCs when compared with *Brca2* hypomorphic mutant ESCs (R2336H), which are defective in HR (*52*) (**Fig. 2C**), suggesting that the double deficient cells were competent for HR.

**Figure 2:**
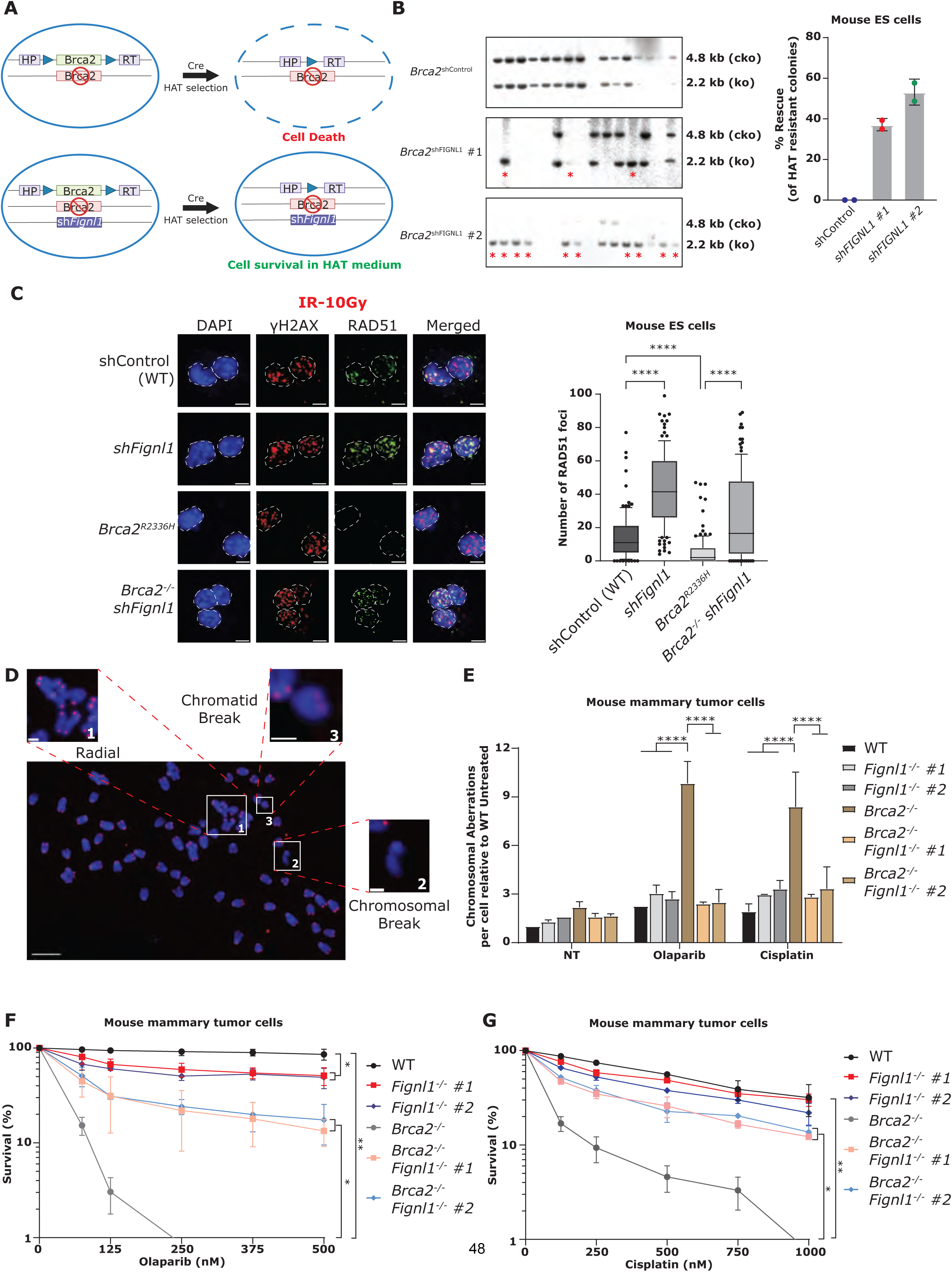
FIGNL1 loss results in synthetic viability of BRCA2 null ESCs and confers resistance to chemotherapeutic drugs. (A) Schematic representation of strategy behind deletion of BRCA2 in mouse embryonic stem cells. (B) Left: representative southern blot images showing the 4.8kb conditional knock-out allele (upper band) and 2.2kb deleted allele (lower band) of BRCA2. Stars in the shFIGNL1 clones represents the viable clones in the absence of BRCA2. Right: Percentage quantification of rescued clones. (C) Left: representative RAD51-IF images in mouse embryonic stem cells (scale bar: 20μm). *Brca2^R2336H^*a Brca2 cell line that has a hypomorphic allele (p.arg2336His) of BRCA2 that is defective in HR. Right: Semi-automated QIBC plots of the RAD51 foci per nucleus in WT and FIGNL1 knock down clones; scatter box plot showing the foci distribution between 10-90th percentile (n=150). The experiment was performed two times. P-values from Mann-Whitney test (two-tailed) comparing between indicated groups (****: <0.0001). (D) Representative metaphase spread image showing different aberrations analyzed for genomic instability. Scale bar (whole metaphase): 5 µm; scale bar (individual aberrations): 1 µm. (E) Quantification of chromosomal aberrations in mouse mammary tumor cells in response to Olaparib and cisplatin from 3 independent experiments in which 50 individual metaphases were analyzed for each sample per experiment. P values from Ordinary two-way ANOVA (****: <0.0001) (F) Percentage survivability of mouse mammary tumor cells in response to Olaparib; quantification was performed by normalization to non-treated cells. P-values from Dunnett’s multiple comparison One-Way ANOVA comparing between indicated groups (*: <0.05, **: <0.01). (G) Percentage survivability of mouse mammary tumor cells in response to cisplatin; quantification was performed by normalization to non-treated cells. P-values from Dunnett’s multiple comparison One-Way ANOVA comparing between indicated groups (*: <0.05, **: <0.01).

Since the loss of FIGNL1 upon BRCA2 deficiency resulted in the restoration of HR, we next assessed the levels of genome instability in these cells in response to chemotherapeutic treatments that induce DSBs. Chromosomal aberrations were quantified by analyzing metaphase spreads upon treatment with the PARP inhibitor (PARPi) olaparib and the interstrand crosslinking agent cisplatin in mouse mammary tumors and RPE1 cells. Interestingly, our data revealed that treatments with either PARPi or cisplatin in BRCA2/ FIGNL1 double deficient cells significantly rescued the levels of chromosomal aberrations that were observed upon loss of BRCA2 alone (**Fig. 2D, E and Fig.S5A**). Similar data were also observed in mESCs, which displayed restoration of genome stability upon PARPi treatment in *Brca2^-/-^ shFignl1* double-deficient cells compared to *Brca2^R2336H^* hypomorphic cells, suggesting that the loss of FIGNL1 under BRCA2-deficient conditions could confer resistance to chemotherapeutic drugs that induce DSBs (**Fig.S5B)**. Finally, to test whether the rescue of genome instability observed in BRCA2/FIGNL1 double-deficient cells also correlates with the increased survival of these cells, we performed clonogenic survival assays in mouse mammary tumors and mESCs. Consistent with our data on genome instability, the loss of FIGNL1 in BRCA2 deficient cells significantly rescued the cellular sensitivity of BRCA2-deficient cells upon treatment with both PARPi and platinum drugs (**Fig.2. F-G and S5 C-F**). Taken together, these data suggest that restoration of functional HR through RAD51 loading in BRCA2 deficient cells drives synthetic viability and chemoresistance upon loss of FIGNL1 expression.

### FIGNL1 interaction with RAD51 is critical for its regulation

FIGNL1 is a multi-domain protein consisting of a FRBD domain that interacts with RAD51, AAA+ ATPase domain, and V domain (*38*) (**Fig.1C and S3H-I**). To identify the region of FIGNL1 involved in the dissociation of RAD51, we expressed FL (full length-FIGNL1), dFRBD (deletion of the FRBD domain that interacts with RAD51), F295E (point mutation in the FxxA motif in the FRBD domain), dNLS (disrupts the interaction with FIRRM/FLIP, which stabilizes FIGNL1), and K447A+D500A (point mutations in the ATPase domain) in *FIGNL1^-/-^* RPE1 cells (**Fig.S6A-B**) and tested for RAD51 focus accumulation upon treatment with IR. As anticipated, reconstitution of FIGNL1-FL resulted in *FIGNL1^-/-^* and shBRCA*2 FIGNL1^-/-^* cells displaying significantly reduced levels of RAD51 foci, similar to the WT or BRCA2 deficient conditions respectively (**Fig.3A**). Strikingly, reconstitution with dFRBD, F295E, dNLS, or K447A+D500A mutant constructs failed to reduce the number of RAD51 foci to WT or BRCA2 deficient levels (**Fig.3A**). Next, to test whether the failure to reduce RAD51 focus levels by the mutants also correlated with genome stability in shBRCA*2 FIGNL1^-/-^* cells, we measured chromosomal aberrations in these cells upon treatment with either PARPi or cisplatin. Reconstitution with FIGNL1-FL and treatment with either PARPi or cisplatin significantly increased chromosomal aberrations in the BRCA2/FIGNL1 double-deficient cells (**Fig.S6C)**. However, consistent with our data on RAD51, the expression of all mutants displayed reduced genome instability, similar to shBRCA*2 FIGNL1^-/-^* cells (**Fig.S6C)**. Taken together, these data suggest that stabilization of FIGNL1 by FIRRM/FLIP and the interaction of RAD51 with both the FRBD and ATPase domains of FIGNL1 are critical for its function in RAD51 dissociation from DSBs.

**Figure 3:**
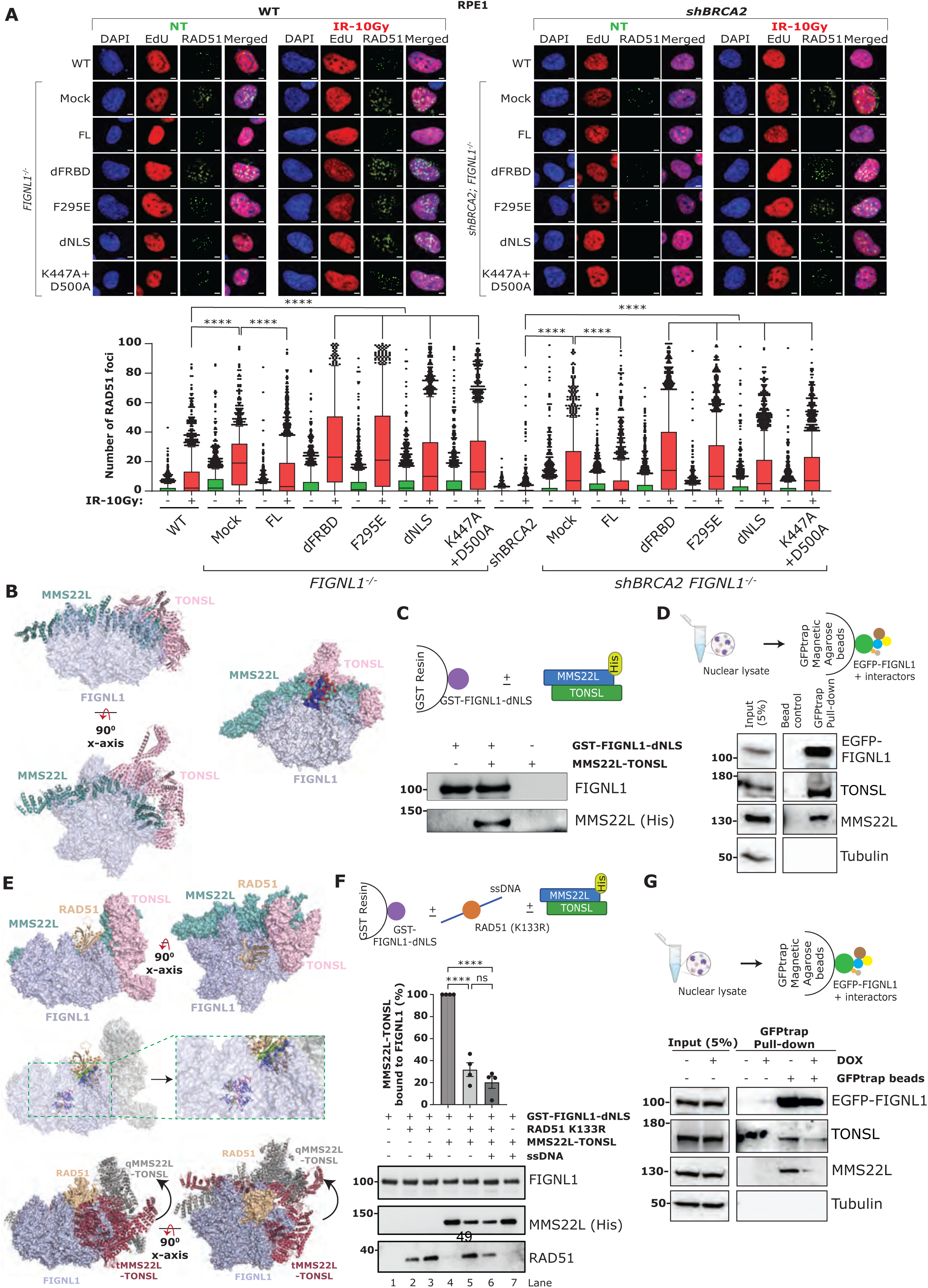
RAD51 availability determines FIGNL1 interaction with MMS22L-TONSL complex via FRBD domain. (A) Top: representative images of high-content, automated RAD51-IF images showing the EdU and RAD51 foci in untreated and ionizing radiation treated human RPE1 cells; scale bar: 5µm. Bottom: Automated QIBC plots of the RAD51 foci per nucleus; scatter box plot showing the foci distribution between 10-90th percentile from 1500 cells (experiment was repeated 3 times). P-values from Mann-Whitney test (two-tailed) comparing between indicated groups (****: <0.0001). (B) Left: The side and top view of FIGNL1-MMS22L-TONSL complex. The FIGNL1 hexamer is shown as surface representation (Light blue) while MMS22L (teal) and TONSL (light pink) are shown in cartoon representation. Right: Surface representation of ternary complex showing the major binding interface of ternary complex. MMS22L-TONSL interacts through its heterodimeric interface (maroon) with FIGNL1 FRBD and NLS-FRBD connecting loop (dark blue). (C) Top: Schematics of the *in vitro* interaction assay between recombinant FIGNL1-dNLS (immobilized) and MMS22L-TONSL. Bottom: Western blot results show that FIGNL1-dNLS interacts with MMS22L-TONSL in the *in vitro* interaction assay. (D) Top: Schematics of the GFPtrap pull-down of the EGFP-FIGNL1 and MMS22L-TONSL from nuclear extracts. EGFP-FIGNL1 stably over-expressing cells in *FIGNL1^-/-^* RPE1 cells were used for nuclear fractionation and GFPtrap pulled down. Bottom: Western blot results indicative of the interaction between FIGNL1 and MMS22L-TONSL. (E) (Top) The side and top of the FIGNL1-RAD51-MMS22L-TONSL complex. The FIGNL1 hexamer, (Light blue), MMS22L (teal), and TONSL (light pink) are shown as surface representation, while RAD51 (beige) is shown in cartoon representation. (Middle) Surface representation and zoom in view of ternary complex showing the binding interfaces of FIGNL1− FRBD and pore loops are represented in dark blue; while RAD51 interacting domains, N-terminal and loop connecting Walker A/B motifs are represented in pink and green respectively. The non-interacting MMS22L-TONSL is marked in grey. (Bottom) Side and top view of the overlay of FIGNL1-MMS22L-TONSL in ternary (raspberry/tMMS22L-TONSL/without RAD51) and quaternary (grey/qMMS22L-TONSL/with RAD51) complexes. The outward movement of the MMS22L-TONSL heterodimeric interface away from the FIGNL1 pocket (shown by black arrow) represents the loss of FIGNL1-MMS22L-TONSL binding upon FIGNL1-RAD51 interaction. (F) Top: Schematics of the *in vitro* interaction assay between recombinant FIGNL1-dNLS (immobilized), MMS22L-TONSL and RAD51-K133R in the presence or absence of ssDNA. Bottom: Percentage normalized quantifications showing the band intensity of the bound MMS22L-TONSL to FIGNL1-dNLS; error bars are indicative of mean±SEM from 4 independent experiments (n=4). P-values from Ordinary One-Way ANOVA comparing between indicated groups (****: <0.0001). Western blot results indicate the interactions between the proteins involved. (G) Top: Schematics of the GFPtrap pull-down of the EGFP-FIGNL1 and MMS22L-TONSL in the presence or absence of BRCA2 (+DOX), from nuclear extracts. EGFP-FIGNL1 stably over-expressing cells in *FIGNL1^-/-^* RPE1 cells were used for nuclear fractionation and GFPtrap pulled down. Bottom: Western blot results indicative of the interaction between FIGNL1 and MMS22L-TONSL.

### FIGNL1 interacts with MMS22L-TONSL complex in the absence of RAD51 availability

The rescue of RAD51 loading and HR in BRCA2/FIGNL1 double-deficient cells indicated the presence of other factors that could mediate RAD51 loading and stimulate HR in the absence of BRCA2. Therefore, to identify factors that could stimulate RAD51-mediated HR and be regulated by FIGNL1, we investigated the potential protein-protein interactions (PPIs) of FIGNL1. To this end, we probed the PREDICTOMES classifier database (https://predictomes.org/view/ddr), which uses AlphaFold multimer to predict potential high-confidence PPIs for DNA replication and repair factors (*53*). Consistent with established data, our search revealed FIRRM, RAD51 and DMC1 as the top hits as potential interactors of FIGNL1 based on <u>p</u>redicted local <u>d</u>istance <u>d</u>ifference test (pLDDT) scores (**Fig.S7A and Table.3**). Interestingly, FIGNL1 was also predicted to interact with high confidence with both MMS22L and TONSL, which exists as a complex and mediates HR though the interaction with RAD51 (**Fig. S7A and Table.3**) (*20–22*). To further analyze the predicted interactions between FIGNL1, MMS22L, and TONSL, we used AlphaFold 3 to model the structures and interactions between FIGNL1, MMS22L, and TONSL in a complex. Assessment of the molecular architecture of the FIGNL1-MMS22L-TONSL complex revealed that majorly two monomeric units of FIGNL1 interact with the MMS22L-TONSL heterodimer **(Fig.3B left, Table.1)**. The dimeric region of MMS22L-TONSL serves as the primary interacting motif for FIGNL1. The FRBD region and FRBD-NLS connecting loop of one FIGNL1 monomer contributed to complex formation via interaction with the MMS22L-TONSL heterodimeric interface (**Fig.3B right, Table.1**). The second FIGNL1 monomer showed a weak interaction with MMS22L. However, no significant binding interactions were predicted between the pore loops (1 and 2) of the FIGNL1 ATPase domain and the MMS22L-TONSL heterodimer. To test the predicted PPI between FIGNL1 and the MMS22l-TONSL complexes, we next performed an *in vitro* interaction assay with purified FIGNL1-dNLS and MMS22L-TONSL complexes (**Fig.S7A and S7B**). Consistent with the Alphafold 3 prediction, the addition of MMS22L-TONSL complex to immobilized FIGNL1-dNLS revealed a direct physical interaction (**Fig.3C)**. To further validate this interaction in cells, we performed immunoprecipitation (IP) experiments using reconstituted EGFP-tagged FIGNL1 in *FIGNL1^-/-^*RPE1 cells and probed for the interaction between FIGNL1, MMS22L, and TONSL. Pulldowns of FIGNL1 from RPE1 cells displayed a clear interaction with both MMS22L and TONSL, confirming our prediction and the *in vitro* experimental data (**Fig.3D)**.

**Table 2:**
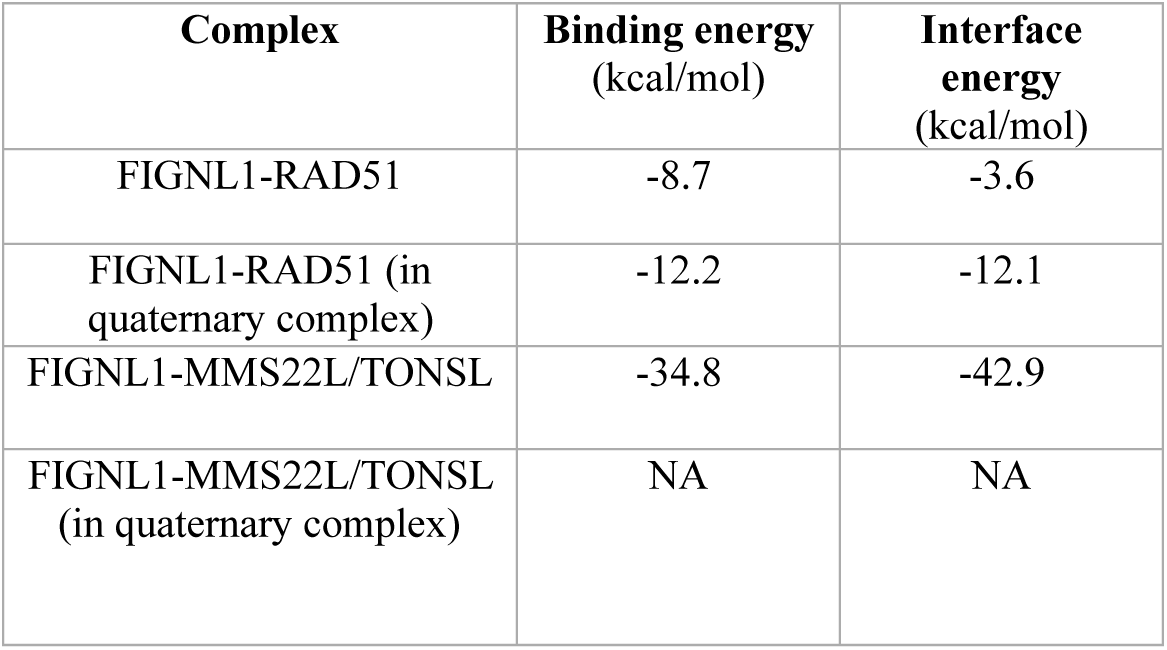
The binding energetics and domains for formation of FIGNL1 complexes with RAD51 and MMS22L/TONSL.

| Complex | Binding energy (kcal/mol) | Interface energy (kcal/mol) |
| --- | --- | --- |
| FIGNL1-RAD51 | -8.7 | -3.6 |
| FIGNL1-RAD51 (in quaternary complex) | -12.2 | -12.1 |
| FIGNL1-MMS22L/TONSL | -34.8 | -42.9 |
| FIGNL1-MMS22L/TONSL (in quaternary complex) | NA | NA |

**Table 3:** Predictome interacting partners of FIGNL1 involved in genome maintenance.

| Sl. No. | Name | pLDDT | Sl. No. | Name | pLDDT | Sl. No. | Name | pLDDT |
| --- | --- | --- | --- | --- | --- | --- | --- | --- |
| 1 | FIRRM | 89.6 | 37 | RMI2 | 72.8 | 73 | REV1 | 67.8 |
| 2 | POLD3 | 87.4 | 38 | TIMELESS | 72.7 | 74 | RAD18 | 67.5 |
| 3 | RAD51 | 86.3 | 39 | RECQL | 72.6 | 75 | PALB2 | 67.5 |
| 4 | CCNA2 | 82 | 40 | DDB1 | 72.5 | 76 | FANCI | 66.7 |
| 5 | DMC1 | 81.7 | 41 | DNA2 | 72.4 | 77 | MCMBP | 66.7 |
| 6 | PCNA | 81.3 | 42 | NEIL1 | 72.4 | 78 | SLF1 | 66.5 |
| 7 | RAD51B | 81.1 | 43 | FANCL | 72.1 | 79 | MAD2L2 | 66.4 |
| 8 | FANCC | 79.6 | 44 | HELQ | 72 | 80 | MUS81 | 66.4 |
| 9 | RAD51C | 79.2 | 45 | AFG2A | 72 | 81 | POLD1 | 66.3 |
| 10 | XRCC3 | 78.9 | 46 | STAG2 | 71.8 | 82 | RPA3 | 65.8 |
| 11 | POLL | 78.9 | 47 | FANCD2 | 71.7 | 83 | BARD1 | 65.5 |
| 12 | FIGNL1 | 78.8 | 48 | PRIM2 | 71.7 | 84 | ETAA1 | 65.2 |
| 13 | CTC1 | 78.7 | 49 | AFG2B | 71.5 | 85 | TP53BP1 | 65.1 |
| 14 | ABRAXAS1 | 77.8 | 50 | PDS5B | 71.5 | 86 | SSRP1 | 65.1 |
| 15 | GINS3 | 77.5 | 51 | ERCC3 | 71.4 | 87 | MCM6 | 64.9 |
| 16 | ESCO1 | 77.5 | 52 | SMC5 | 71 | 88 | RNASEH2C | 64.5 |
| 17 | CHTF8 | 76.8 | 53 | RAD52 | 70.9 | 89 | PDS5A | 64.2 |
| 18 | RFC2 | 76.7 | 54 | MMS22L | 70.7 | 90 | PRIMPOL | 63.7 |
| 19 | ORC5 | 76.1 | 55 | TOP1 | 70.7 | 91 | RFC3 | 63.5 |
| 20 | POLA2 | 76 | 56 | UBE2T | 70.5 | 92 | XPC | 63.5 |
| 21 | EXO5 | 75.7 | 57 | MCM9 | 70.1 | 93 | WAPL | 62 |
| 22 | RAD51D | 75 | 58 | SUPT16H | 70 | 94 | ATR | 61.5 |
| 23 | SHLD3 | 75 | 59 | GINS1 | 69.9 | 95 | MCM7 | 61.4 |
| 24 | TERF1 | 74.7 | 60 | UBE2B | 69.9 | 96 | MSH6 | 60.1 |
| 25 | DTL | 74.5 | 61 | CDK2 | 69.5 | 97 | DCLRE1A | 59.7 |
| 26 | WRAP53 | 74.5 | 62 | CDT1 | 69.4 | 98 | EME1 | 58.9 |
| 27 | STN1 | 74.1 | 63 | SETMAR | 69.4 | 99 | GMNN | 58.5 |
| 28 | GINS2 | 74 | 64 | ERCC4 | 69.1 | 100 | TICRR | 57.4 |
| 29 | MAU2 | 74 | 65 | RECQL4 | 69 | 101 | SLX4 | 56.9 |
| 30 | ORC6 | 73.9 | 66 | FAAP100 | 68.7 | 102 | MLH3 | 55.8 |
| 31 | ORC1 | 73.9 | 67 | RFC1 | 68.6 | 103 | FAAP20 | 54.8 |
| 32 | BLM | 73.3 | 68 | TDP2 | 68.5 | 104 | FAN1 | 54.1 |
| 33 | TIPIN | 73.3 | 69 | TONSL | 68.4 | 105 | TTF2 | 53.4 |
| 34 | CHTF18 | 73.1 | 70 | DBF4B | 68.3 | 106 | RNASEH2B | 52.5 |
| 35 | TERF2 | 73.1 | 71 | MSH3 | 68.2 |  |  |  |
| 36 | CHAF1B | 72.8 | 72 | POLE | 67.8 |  |  |  |

Since both the MMS22L-TONSL complex and FIGNL1 modulate RAD51 activity in HR, we next predicted the interactions of the FIGNL1-MMS22L-TONSL ternary complex in the presence of RAD51 using AlphaFold 3. In the quaternary complex FIGNL1-RAD51-MMS22L-TONSL **(Fig. 3E top)**, as expected RAD51 was predicted to bind to FIGNL1 in the same structural pocket as that observed previously in the absence of MMS22L-TONSL complex **(Fig. 1C)**. Interestingly however, RAD51 was observed to be in a tilted orientation and exhibited interactions with the FxxA and FxxP regions of FIGNL1-FRBD pocket **(Fig.3E middle)**. Interface and binding energy calculations of the dimeric and quaternary structures of FIGNL1-RAD51 predicted that, in both cases, RAD51 was highly stabilized in its FIGNL1 binding pocket **(Table.2)**. Strikingly, the addition of RAD51 to the FIGNL1-MMS22L-TONSL ternary complex predicted an outward shift of the MMS22L-TONSL dimeric interface domain from FIGNL1, suggesting the dissociation of the MMS22L-TONSL heterodimer from the FIGNL1 hexamer **(Fig.3E bottom)**. This dissociation is also evident from the calculations of the binding and interface energy parameters, where no significant favourable energies were predicted in the quaternary complex, which is in sharp contrast to the ternary complex **(Table.2)**.

To further test this prediction of dissociation of the MMS22L-TONSL complex from FIGNL1 in the presence of RAD51, we performed *in vitro* interaction assays with purified FIGNL1-dNLS and MMS22L-TONSL complex in the presence or absence of RAD51 and/or ssDNA-bound RAD51. To ensure stronger binding of RAD51 to ssDNA, RAD51-K133R (a mutant defective in ATP hydrolysis) (*54, 55*) was used in the reactions (**Fig.S7B-C)**. Consistent with our structural prediction data, the presence of RAD51-K133R significantly reduced the interaction between the MMS22L-TONSL complex and FIGNL1. Interestingly, no further significant decrease in interaction was observed upon the addition of ssDNA-bound RAD51 to the reaction when compared to RAD51 alone **(Fig.3F)**.

BRCA2 has been proposed to sequester most cellular RAD51 to prevent inappropriate DNA interactions (*56*). Therefore, we hypothesized that RAD51 availability for the MMS22L-TONSL-FIGNL1 complex in cells could be constrained in the presence of BRCA2. To test this hypothesis, we performed pull-down experiments for FIGNL1 in the presence or absence of BRCA2 in RPE1 cells. Strikingly, our data showed that BRCA2 deficiency resulted in the dissociation of the interaction between FIGNL1 and the MMS22L-TONSL complex when compared to WT cells **(Fig.3G)**. Taken together, these data strongly suggest that the MMS22L-TONSL-FIGNL1 interaction may be critical for controlling FIGNL1 activity and limiting unwanted RAD51 removal during HR.

### MMS22L-TONSL promotes RAD51 accumulation and facilitates HR in BRCA2/FIGNL1 double-deficient cells

Our data suggest that the interaction between MMS22L-TONSL and FIGNL1 is important for the regulation of RAD51 dynamics upon DSB induction. To test this, we assessed the localization of MMS22L to DSBs in the presence or absence of BRCA2 and/or FIGNL1 in RPE1 and mouse mammary tumor cells. Upon exposure to IR, WT cells displayed significantly increased MMS22L localization compared to untreated (NT) cells **(Fig.4A and S8A)**. Interestingly, the loss of FIGNL1 in two separate clones resulted in a significant increase in MMS22L foci compared to WT cells. Loss of BRCA2 resulted in a slight but significant increase in MMS22L foci numbers, which was significantly enhanced in BRCA2/FIGNL1 double-deficient cells **(Fig.4A and S8A)**. This increase in MMS22L localization to DSBs upon loss of FIGNL1 in BRCA2 proficient or deficient conditions was also highly correlated with increased RAD51 localization to DSBs upon IR exposure (**Fig.1A and S2A**). Taken together, these data suggest that FIGNL1 primarily dissociates RAD51 loading mediated by a pathway comprising the MMS22L-TONSL complex to fine-tune HR efficiency. To validate this hypothesis directly, we performed *in vitro* DNA strand exchange reactions in the presence of the purified MMS22L-TONSL complex, RAD51, and FIGNL1-dNLS **(Fig.4B)**. MMS22L-TONSL complex was observed to significantly stimulate the strand exchange mediated by RAD51 consistently with previous evidence (*22*). However, upon the addition of FIGNL1-dNLS, the strand exchange reaction was notably inhibited in a concentration-dependent manner **(Fig.4B)**. These data strongly implicate that, unlike BRCA2, the MMS22L-TONSL complex is unable to counteract FIGNL1 mediated RAD51 dissociation, thus resulting in the inhibition of HR.

**Figure 4:**
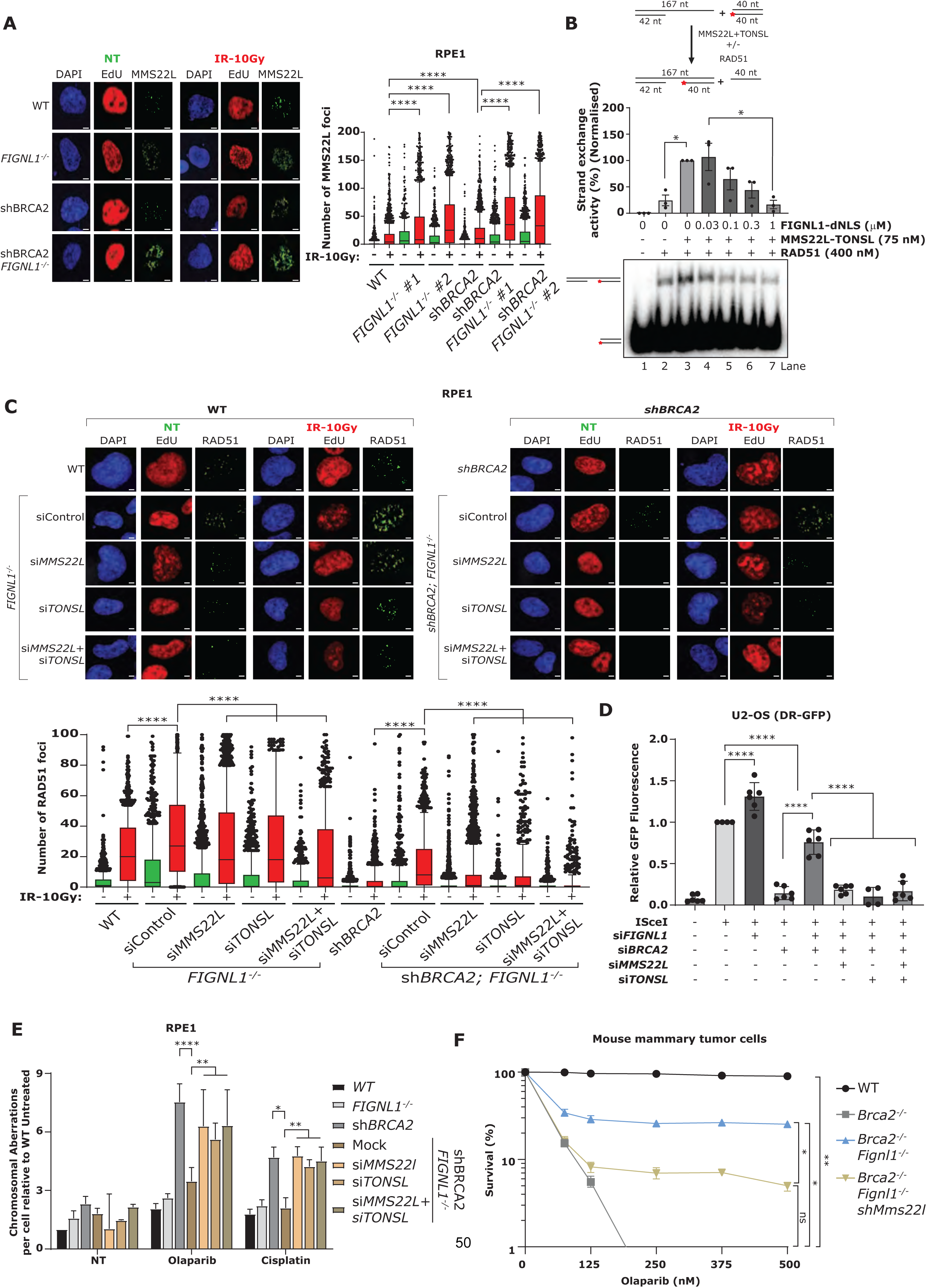
MMS22L-TONSL complex loads RAD51 in the BRCA2-FIGNL1 deficient cells to recue HR and confer chemoresistance. (A) Left: Representative high-content, automated IF images showing EdU and MMS22L foci in untreated and ionizing radiation treated cells; scale bar: 5µm. Right: Automated QIBC plots of the MMS22L foci per nucleus; scatter box plot showing the foci distribution between 10-90th percentile from 2000 cells (experiment was repeated 3 times). P-values from Mann-Whitney test (two-tailed) comparing between indicated groups (****: <0.0001). (B) Top: Schematics of the DNA strand exchange assay. The strand exchange assays were performed by adding 400nM purified RAD51 and 75nM MMS22L-TONSL to the DNA substrate. The assay was then supplemented by increasing concentrations of purified FIGNL1-dNLS. The red asterisk indicates the position of the radioactive label. Bottom: Percentage normalized quantifications showing the band intensity of the exchanged DNA; error bars are indicative of mean±SEM from 3 independent experiments (n=3). P-values from ordinary One-way ANOVA comparing between indicated groups (*: <0.05). (C) Top: Representative high-content, automated IF images showing EdU and RAD51 foci in untreated and ionizing radiation treated human RPE1 cells; scale bar: 5µm. Bottom: Automated QIBC plots of the RAD51 foci per nucleus; scatter box plot showing the foci distribution between 10-90th percentile from 1500 cells (experiment was repeated 3 times). P-values from Mann-Whitney test (two-tailed) comparing between indicated groups (****: <0.0001). (D) DR-GFP system stably integrated U2-OS cells transfected with 72 hours of indicated siRNAs overlapping with 48 hours of ISceI to measure the relative levels of GFP fluorescence to the WT cells. Error bars are representative of the mean±SD from 3 independent experiments (n=3). P-values from unpaired t-test comparing between indicated groups (****: <0.0001). (E) Quantification of chromosomal aberrations in RPE1 FIGNL1 knock-out clones with siRNAs against MMS22L and TONSL in response to Olaparib and cisplatin from 3 independent experiments in which 50 individual metaphases were analyzed for each sample per experiment. P values from Ordinary two-way ANOVA (*: <0.05; **: <0.01; ****: <0.0001). (F) Percentage survivability of mouse mammary tumor cells in response to Olaparib; quantification was performed by normalization to non-treated cells. P-values from Dunnett’s multiple comparison One-Way ANOVA comparing between indicated groups (ns: 0.1234; *: <0.05, **: <0.01).

Because MMS22L localization was enhanced upon the loss of BRCA2/FIGNL1 double-deficient cells, we tested whether the MMS22L-TONSL complex indeed mediates the rescue of RAD51 loading at DSBs in these cells. To this end, we downregulated MMS22L, TONSL, or both in *FIGNL1^-/-^* and *shBRCA2 FIGNL1^-/-^* RPE1 cells (**Fig. S8B**) and probed for RAD51 accumulation upon induction of DSBs. Interestingly, downregulation of the MMS22L-TONSL complex significantly decreased RAD51 foci accumulation in *FIGNL1^-/-^*cells to nearly WT levels, again supporting the notion that FIGNL1 primarily dissociates RAD51 accumulation mediated by the MMS22L-TONSL complex **(Fig.4C)**. Strikingly however, loss of the MMS22L-TONSL complex in BRCA2/FIGNL1 double-deficient cells almost completely abrogated RAD51 foci accumulation. These data strongly suggest that the MMS22L-TONSL complex is essential for RAD51 accumulation at DSBs and restoration of HR in BRCA2/FIGNL1 double-deficient cells **(Fig.4C)**. Next, we assessed HR efficiency by performing DR-GFP reporter assays upon the loss of the MMS22L-TONSL complex in BRCA2/FIGNL1 double-deficient cells. In agreement with our data on RAD51 accumulation, loss of the MMS22L-TONSL complex in BRCA2/FIGNL1 double-deficient cells resulted in significantly reduced HR efficiency compared to BRCA2/FIGNL1 double-deficient cells alone **(Fig.4D and S8C)**. Taken together, these results confirm the critical role of MMS22L-TONSL in mediating HR upon the loss of BRCA2 and FIGNL1 expression. Next, we tested whether loss of MMS22L-TONSL mediated HR defects in BRCA2/FIGNL1 double-deficient cells renders them genomically unstable upon treatment with DSB-inducing chemotherapeutic treatments. Therefore, we analyzed metaphase spreads to assess chromosomal aberrations in RPE1 cells following siRNA-mediated downregulation of the MMS22L-TONSL complex. Our data revealed that loss of the MMS22L-TONSL complex in sh*BRCA2 FIGNL1^-/-^*cells displayed significantly increased levels of chromosomal aberrations upon treatment with both PARPi and cisplatin when compared to *shBRCA2 FIGNL1^-/-^* cells **(Fig.4E)**. Finally, we tested whether the genome instability observed upon the loss of MMS22L-TONSL also correlated with cellular sensitivity to chemotherapeutic drugs. Therefore, we performed clonogenic survival assays upon treatment with PARPi by inducibly downregulating MMS22L in *Brca2^-/-^ Fignl^-/-^* mouse mammary tumor cells (**Fig.S8D**). In line with the genome instability data, our data revealed that the loss of MMS22L in *Brca2^-/-^ Fignl^-/-^* cells resulted in significant cellular sensitivity to PARPi treatment when compared *to Brca2^-/-^ Fignl^-/-^* cells alone **(Fig.4F)**. Taken together, these data underscore the essential role of MMS22L-TONSL mediated stimulation of HR to maintain genome stability in BRCA2- and FIGNL1-deficient cells.

## DISCUSSION

The loss of RAD51 nucleation and stabilization is considered to be the primary cause of HR defects and loss of cellular viability upon BRCA2 deficiency (*5–11*). There have been no reports to date demonstrating compensatory pathways that can restore RAD51 loading and HR function in the absence of BRCA2. This study presents the first instance of a genetic alteration in mammalian cells, wherein the loss of the AAA+ ATPase FIGNL1 restores RAD51 loading, functional HR, and genome stability in BRCA2 deficient cells. This restoration of HR rescues the lethality in mESCs and confers PARPi and cisplatin resistance in *Brca2*-null cells. The observed restoration of RAD51 loading and subsequent HR in BRCA2-deficient cells indicates that the HR defects observed upon BRCA2 loss cannot be attributed solely to the lack of RAD51 loading and stabilization, but rather to the unrestricted dissociation of RAD51 from the resected ends by FIGNL1 (**Fig.5**). Furthermore, our data indicate that fine-tuning RAD51 levels at DSBs through both loading and dissociation is crucial for HR fidelity and efficacy (**Fig.5**). In accord, both increased and decreased RAD51 levels are incompatible with cellular viability (*9, 40, 42, 50*).

**Figure 5:**
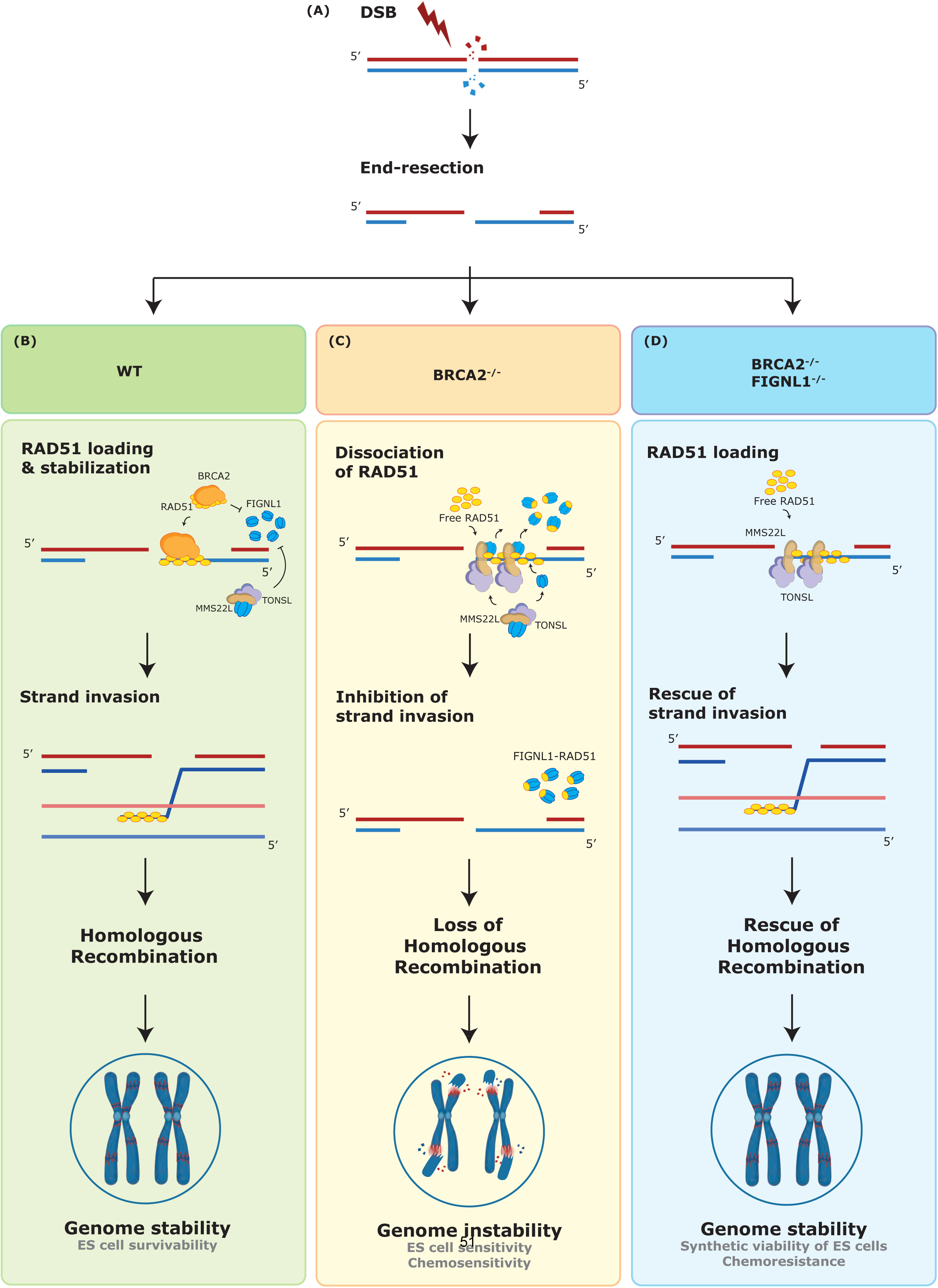
Proposed model of the study. (A) Upon induction of DSBs, the break ends are resected in the 5’to 3’ direction, exposing a 3’ssDNA overhang. (B) BRCA2 plays a key role by loading and stabilizing RAD51 on the resected single-stranded ssDNA, facilitating RAD51-mediated strand invasion. Besides facilitating RAD51 loading, BRCA2 is crucial for maintaining RAD51 filament stability by preventing its premature dissociation caused by the anti-recombinase FIGNL1. In the presence of BRCA2, the limited availability of free RAD51 allows the hexameric FIGNL1 to interact with the MMS22L-TONSL complex. This interaction ensures that RAD51 levels are finely regulated during HR in wild-type cells. (C) However, in the absence of BRCA2, the elevated concentration of free RAD51 results in the dissociation of the FIGNL1-MMS22L-TONSL complex. This could lead to in dynamic cycles of RAD51 loading and dissociation between the MMS22L-TONSL complex and FIGNL1, ultimately causing RAD51 loading defect and increased genome instability. (D) The loss of FIGNL1 in BRCA2-deficient cells restores productive RAD51 loading by the MMS22L-TONSL complex, rescuing HR efficiency, which in turn restores the genome instability observed upon loss of BRCA2.

Our biochemical data suggest that FIGNL1 can directly dissociate RAD51 from ssDNA and inhibit DNA strand invasion. In contrast to other known anti-recombinases (*27–32*), FIGNL1 affinity to DNA is very low, and hence its function is likely dependent on direct physical interactions rather than primarily DNA binding. Consistent with a recent report from the Jasin and Zhang labs (*42*), our structure prediction model shows that FIGNL1 interacts with RAD51 through the FRBD domain and the Pore Loops, which are embedded in the ATPase domain of FIGNL1. In line with this prediction, reconstitution of *FIGNL1^-/-^* cells with deletions and mutations within the FRBD and ATPase domains results in retention of high RAD51 foci and also rescues the RAD51 foci formation defects upon BRCA2 deficiency. Interestingly, our cellular and biochemical data strongly suggest that the presence of BRCA2 can inhibit the dissociation of RAD51 by FIGNL1 to allow strand exchange. This inhibition of FIGNL1 activity by BRCA2 could be due to its high-affinity binding and sequestration of the majority of nuclear RAD51, as well as its role in enhancing RAD51 stability on ssDNA thus preventing RAD51 dissociation (*3, 5, 56, 57*). The inhibition of FIGNL1 activity by BRCA2 may serve as a regulatory mechanism to prevent excessive dissociation of RAD51 from DSBs to ensure the fidelity of the HR process (**Fig.5**).

Absence of FIGNL1 in BRCA2 proficient cells leads to a significant increase in RAD51 accumulation at DNA damage sites. This suggests that FIGNL1’s primary function is to dissociate RAD51 that has been loaded by an unknown mediator. Our data show that FIGNL1 interacts with the MMS22L-TONSL complex, which has previously been shown to stimulate RAD51 loading at DSBs and promote DNA strand exchange (*20–22*). Furthermore, loss of FIGNL1 showed significantly higher localization of MMS22L to DSBs suggesting that FIGNL1 dissociates RAD51 loading that is dependent on the MMS22L-TONSL complex, possibly in association with additional factors (**Fig.5**). Consistent with this idea, we demonstrated FIGNL1 to have the ability to effectively suppress the strand exchange reaction facilitated by the MMS22L-TONSL complex. Notably, the association between the MMS22L-TONSL complex and FIGNL1 was found to be disrupted under two conditions: *in vitro* when RAD51 was present, and in cells lacking BRCA2, where RAD51 is unbound to BRCA2 and available.

We speculate that the interaction between the FIGNL1 and MMS22L-TONSL complex may represent an additional layer of regulation to fine-tune the fidelity of HR. In this scenario, in addition to being inhibited by BRCA2, FIGNL1 activity could be further fine-tuned via its interaction with MMS22L-TONSL into a ternary complex when RAD51 availability is limiting in BRCA2 proficient conditions (**Fig.5**). In line with this concept, our Alphafold predictions show that RAD51 binding FRBD region interacts with the MMS22L-TONSL complex in the absence of RAD51. However, in the presence of RAD51, this complex is disrupted, resulting in an energetically favorable interaction between RAD51 and the FRBD region of FIGNL1. This situation might represent a BRCA2-deficient state where RAD51 is readily accessible, thus facilitating the removal of RAD51 from DSBs by FIGNL1, which is initially loaded by the MMS22L-TONSL complex and possibly additional co-factors (**Fig.5**).

Our findings align with a model where the MMS22L-TONSL complex facilitates HR in the absence of BRCA2 and FIGNL1 (**Fig.5**). The association of TONSL with H4K20me0, along with its interaction with ASF1 and CAF1, may potentially mediate the recruitment of MMS22L-bound RAD51 to sites of DSBs on newly replicated DNA, thereby facilitating RAD51 loading in BRCA2/FIGNL1-deficient cells (*23, 24*). Supporting this concept, the loss of the MMS22L-TONSL complex in cells lacking both BRCA2 and FIGNL1 led to an abrogation in RAD51 accumulation and impaired HR at DSBs. Additionally, loss of MMS22L-TONSL complex in cells deficient in both BRCA2 and FIGNL1 increased genomic instability and heightened sensitivity to PARPi. This suggests that BRCA2-deficient cells rely entirely on MMS22L-TONSL-mediated HR for survival when challenged by DSBs in the absence of FIGNL1 (**Fig.5**).

Our study has also implications for the understanding of chemoresistance mechanisms in BRCA2-mutated tumors that restore RAD51 loading and HR without reversion mutations in BRCA2 gene during the course of therapy (*13, 58*). In summary, our findings demonstrate that inhibiting RAD51 dissociation from DSBs in BRCA2-deficient cells promotes synthetic viability and drug resistance by restoring HR. Targeting alternate RAD51 loading pathways including MMS22L-TONSL complex could be an attractive strategy to re-sensitize these resistant tumors highlighting the crucial role of precisely regulating RAD51 dynamics during the HR process.

## ACKNOWLEDGEMENTS

The authors thank Optical Imaging Center of the Erasmus MC for assistance and microscope access for the study. The authors thank Dr. Maria Jasin (Memorial Sloan Kettering Cancer Center) for the miBRCA2 expression construct and Prof. Wim Vermeulen for critical reading of the manuscript and discussions

## Author Contributions

*Conceptualization*: A.R.C and Rp.K. *Methodology*: Rp.K, A.A, S.K Sengodan, C.F, N.N, A.FT. *Investigation*: Rp.K, A.A, S.K Sengodan, C.F, N.N, O.I, S.N, E.M.M, K.dK. *Funding acquisition:* A.R.C, P.C, S.K.Sharan, K.M.P, J.H.G.L, R.K. A.FT. Project administration A.R.C. *Supervision:* A.R.C, P.C, S.K.Sharan, K.M.P, J.H.G.L, R.K. *Writing-original draft:* ARC with contributions from Rp.K and CF. *Writing-review and editing:* P.C, S.K.Sharan, K.M.P, J.H.G.L, R.K. and all other authors.

## MATERIALS AND METHODS

### Cell Culture

Human RPE1-hTERT *TP53^-/-^; shBRCA2* cells were cultured in DMEM/F12+GlutaMAX^TM^ (GIBCO) media containing 10% FBS (Capricorn) and 1% penicillin-streptomycin (PS) (GIBCO) at 37°C and 5% CO_2_ in a humidified incubator. Mouse mammary tumor cells KB2PR and KB2P cells (previously described(*59*)), were cultured in DMEM (Gibco) media supplemented with 10% FBS, 1% PS, 5µg/mL insulin (Merck), 5ng/ml murine epidermal growth factor (Merck) and 5 ng/ml cholera toxin and grown under low oxygen conditions (3% O_2_, 5% CO_2_ at 37°C). HEK293T, BOSC and U2-OS (both DIvA(*60*) and DR-GFP(*61*)) cells were cultured in DMEM supplemented with 10% FBS, 1% PS, 1x Non-Essential Amino Acids Solution (Gibco) and 1% Glutamax^TM^ (Gibco) at 37°C and 5% CO_2_ in a humidified incubator. All the cell lines were authenticated by short tandem repeat profiling and routinely tested for mycoplasma contamination.

### Cloning and generation of stable cell line models

Human FIGNL1 sgRNAs (table below) were cloned into pDG458 (Addgene #100900) plasmid at BbsI site on both side of Cas9 expression plasmid. Mouse Fignl1 sgRNA start and stop sequences (table below) were cloned into pSpCas9(BB)-2A-GFP (pX458) (Addgene #48138) at BbsI site. To generate FIGNL1 knock-out models in both human RPE1 and mouse mammary tumor cells, sgRNA cloned plasmids were transfected and FACS sorted for GFP positive cells. Sorted GFP positive cells were then seeded as single cell per well in 96 well plate. Clones grown from single cells were then genotyped and analyzed by western blot to confirm homozygous knockouts.

FIGNL1 mutant constructs: full-length FIGNL1, RAD51 interaction mutants dFRBD (Δ295-344) and FxxA point mutant (F295E); FIRRM/FLIP interaction mutant: dNLS (Δ1-120); and ATPase point mutant (K447A+D500A) cloned in entry clones were used for LR reaction to generate expression clones using Gateway cloning (*35*). Gateway clonase LR enzyme mix II (Invitrogen) was used to clone FIGNL1 mutants into destination vector pCW57.1-EGFP vector (pCW57.1-EGFPnls for dNLS mutant). To create vector backbones for FIGNL1 mutants, Tetracycline-Responsive promoter from pCW57.1-TRE-EGFP/EGFPnls vector were replaced with CMV promoter to create pCW57.1-CMV-EGFP/EGFPnls backbone using NEBuilder® HiFi DNA Assembly Master Mix. Finally, FIGNL1 mutant constructs were co-transfected into HEK293T cells with 3^rd^ generation lentiviral packaging and envelop vectors to create viral particles and transduced into RPE1 FIGNL1 knock-out cells. Transduced RPE1 FIGNL1-KO cells were then sorted using GFP FACS and analyzed for EGFP fused FIGNL1 fusion protein using western blot analysis.

KB2P-AsiSI cells were generated by transfecting doxycycline-inducible AsiSI-YFP plasmid pTRE3G-HA-ER-AsiSI-PGK-GFPVenus-IRES-rtTA3 (a kind gift from Andres Canela, Kyoto University, Japan) into BOSC cells along with pCL-Eco retro-viral packaging vector. Retro-viral particles were collected after 48 hours and transduced into KB2P cells. Transduced cells were then FACS sorted for YFP and clones were selected with equal number of AsiSI induced damage marked by 53BP1 foci levels.

For recombinant FIGNL1-dNLS protein purification, cDNA sequence of FIGNL1-dNLS was cloned into pFB-2xMBP-10xHis Baculovirus backbone or pGEX-2T backbone using NEBuilder® HiFi DNA Assembly Master Mix.

The miBRCA2 construct was PCR amplified from the plasmid pCAGGS_Mini_BRCA2-1xFlag(*62*) (kind gift from Maria Jasin) without the Nuclear Localisation Signal (NLS) using the primers Mini_BRCA2_F and Midi1_BRCA2_BRC1-2_NotI_R with introducing NheI and NotI site at the ends. The product is then cloned into the vector pFastBac-2XMBP-HLTFco-His (*63*) between the NheI and NotI sites to generate the final plasmid pFB-2xMBP-Mini_BRCA2-FLAG-His. All the primers used in the study are listed in the table below.

### Transfection and Infection

All the plasmid DNA transfections were performed using X-tremeGENE™ 9 DNA Transfection Reagent (Merck) and siRNAs were transfected using Lipofectamine™ RNAiMAX Transfection Reagent (Invitrogen) according to manufacturer’s instructions. For viral infections, 48-hour viral soups collected from HEK293T (lenti-viral) and BOSC (retro-viral) cells were transduced into target cell lines with 8µg/ml polybrene (Sigma-Aldrich). Viral titers were assessed by qPCR Lenti/Retrovirus Titer Kits (ABM, Canada).

### ES cell culture and generation of stable lines

PL2F7 mouse embryonic stem cells (mESCs) used in the study were generated from AB2.2 mESC line by knocking out one copy of *Brca2* and flanking the other allele with two *loxP* sites along with 5’ and 3’ human *HPRT minigene*(*64*). mESCs were cultured on mitotically inactive SNL feeder cells in M15 (knockout DMEM) media (Life technologies) as described previously (*64*).

For generation of stable *Fignl1* knockdown in PL2F7 mESC, two different shRNAs against *Fignl1* (table below) and one control shRNA were cloned into PLKO lentiviral vector. Sh*Fignl1* lentiviral particles were generated by co-transfecting HEK293T cells with PLKO_sh*Fignl1* plasmid and packaging plasmid for 48 h. PL2F7-A10 (PL2F7 clone lacking puromycin resistance gene) mESCs were transduced with PLKO_sh*Fignl1* or PLKO_Control lentiviral particles on feeder free gelatinized plate. After 48 h, cells were trypisized and 10^4^ cells were seeded on a 10cm plate containing SNL feeder cells. Cells were selected with puromycin (3ug/ml) for 5 days. Visible colonies were picked and the stable knockdown of sh*Fignl1* were confirmed by western blot.

### Deletion of *Brca2 cko* allele and confirmation of sh*Fignl1 Brca2^ko/ko^* mESCs by Southern blot

sh*Fignl1* expressing PL2F7-A10 stable clones were used for deletion of *Brca2 cko* allele. Ten million mESCs were electroporated with 25ug of *Pgk-Cre* plasmid. After 36h of electroporation, recombinant clones were selected in HAT media for 5 days followed by HT selection for 2 days. Cells were then maintained in M15 media, and mESC colonies were picked into 96 well plate for expansion and genomic DNA isolation using mESCs lysis buffer containing 10mg/ml of proteinaseK as described previously (*64*). Genomic DNA was digested with *EcoRV* at 37°C overnight and the digested DNA was electrophoresed on 1% agarose gel in 1X TBE buffer, transferred to Hybond-N+ nylon membrane. DNA probe was labelled with [αP^32^]-dCTP, hybridized overnight, washed and imaged using Amersham Typhoon image scanner (Cytiva) as described previously (*64*). The PCR based genotyping was performed as described previously (*19*) to detect the *Brca2 cko* and *Brca2 ko* alleles, using the primers listed in the table below.

### Western Blot

Cells for western blot were harvested and lysed in lysis buffer [50mM Tris-HCl (pH 8.0), 150mM NaCl, 0.1% Triton X-100, 0.5% sodium deoxycholate, 0.1% SDS, 2mM EDTA and protease inhibitor cocktail (PIC)]. The protein concentration was determined using the Pierce BCA protein Assay kit. Proteins were loaded and run on a 4-12% Bis-Tris pre-cast gel or a 3-8% Tris-Acetate pre-cast gel (Invitrogen) and were electrophoretically transferred to a PVDF membrane (Millipore). After transfer, the membranes were blocked in 5% milk for 1 hour and incubated overnight in primary antibodies (table below). Membranes were then washed four times in 0.1% TBST and probed with horseradish peroxidase-conjugated secondary antibodies (Cytiva). Finally, the ECL Prime Western Blotting Detection Reagent kit (GE Healthcare) was used to develop the blot.

### Flow-cytometry analysis of cell cycle

For flow-cytometry analysis of cell cycle, the cells were labeled with EdU for 30 minutes followed by fixation in 4% formaldehyde/PBS ant permeabilization in 1% saponin buffer in 1% BSA/PBS. The Click-It reaction was performed using Fluor Azide 594 according to the manufacturer’s instructions (Invitrogen) to label the EdU. The cells were treated with DAPI as a nuclear stain. The samples were measured in BD LSR Fortessa and analyzed by FlowJo software v10.5.0.

### High Content Microscopy

Immunofluorescence staining was performed as described previously (*65*). Briefly, the RPE1 and mouse mammary tumor cells were seed in 96- or 364-well plates. 48 hours (RPE1 cells) or 24 hours (mouse mammary tumor cells) after seeding the cells were treated with EdU for 20 minutes before irradiation with 10Gy of X-ray. Cells were maintained at 37°C for 4 hours before the cells were pre-extracted with ice-cold CSK buffer [10mM PIPES (pH 6.8), 100mM NaCl, 300mM sucrose, 1mM EGTA, 1mM MgCl_2_, 1mM DTT and PIC], fixed with 4% formaldehyde and permeabilized with 0,1% Triton X-100. The cells were blocked in 5% BSA/PBS before they underwent the Click-IT reaction with Alexa Fluor 488 (Invitrogen) according to the manufacturer’s instruction. The cells were washed and then incubated with corresponding primary antibodies for 1 hour 30 minutes at room temperature. Next, they were incubated with Alexa Fluor-conjugated secondary antibodies (Invirogen) for 1 hour at room temperature, after which the cells were treated with DAPI as a nuclear stain. The images were captured with the Opera Phenix High-Content spinning disc confocal microscope and analyzed using Cell Profiler.

For AsiSI induced DSB-IF in KB2P-AsiSI cells, HA-tagged AsiSI was expressed by adding 2µg/ml of doxycycline for 24 hours with last 5 hours of 400nM 4OHT for nuclear translocation. While DivA-AsiSI cells with constitutive AsiSI expression were treated with 400nM 4OHT after 24 hours of seeding. Cells were then processed similarly as mentioned above.

mESCs were seeded on a 24-well plate and after 24h, cells were irradiated at 10Gy. Cells were maintained at 37^°^C for 4h in fresh M15 media before it was fixed and stained for RAD51. Imaging was performed using Confocal spinning Disk microscopy (Opera Phenix) at 40x maginfication and maximum intensity projection were used to count RAD51 foci. DAPI was used as a nuclear stain.

### Automated image analysis

Z-stack images (3 stacks of 1µm height) of individual channels from high throughput microscopy images were merged to create maximum intensity projection and subjected to Cell Profiler analysis. Briefly, nuclei were identified using the DAPI channel and used as mask to identify and measure the intensities of other corresponding channels. Foci detection was performed from masked nuclear regions in the corresponding channels based on foci’s local intensity maxima to the background. For unbiased detection of foci, a single threshold was set for all the groups and exported the quantification to the TIBCO Spotfire software/Graphpad Prism for visualization.

### Structural analysis of FIGNL1 molecular complexes

The full-length amino acid sequences for FIGNL1 (Q6PIW4), RAD51 (Q06609), MMS22L (Q6ZRQ5), TONSL (Q96HA7) were obtained from UniProt. Structures of FIGNL1 hexamer and its complexes with RAD51, MMS22L/TONSL for formation of binary (FIGNL1-RAD51), ternary (FIGNL1-MMS22L/TONSL), and quaternary (FIGNL1-RAD51-MMS22L/TONSL) were generated by Alphafold3(*66*). As Alphafold3 has a cutoff value of 5000 residues for structure prediction, the initial ternary/quaternary complexes were generated by deleting the amino acid sequences of terminal/non-interacting domains of the respective interacting proteins. Once the ternary/quaternary complexes were generated, the deleted terminal regions of FIGNL1, MMS22L, and TONSL were stitched back using *pair_fit* module of PyMol 2.2.3 for generation of full-length protein complexes (https://legacy.ccp4.ac.uk/). Further, to remove steric clashes present in FIGNL1 hexamer/binary/ternary and quaternary structures, these complexes were energy minimized with the steepest descent algorithm using AMBERff99 force field embedded in UCSF Chimera (*67*). The interacting domains of FIGNL1 hexamer, FIGNL1-RAD51, FIGNL1-MMS22L/TONSL, and FIGNL1-RAD51-MMS22L/TONSL complexes were evaluated with the help of PRODIGY (*68*) and PDB-PISA webserver (https://www.ebi.ac.uk/pdbe/pisa/). PRODIGY and PDB-PISA were also applied respectively to calculate the binding energies of the complexes and the interface binding energies of the interacting partners. In case of binary, ternary, and quaternary complexes, hexameric FIGNL1, MMS22L/TONSL heterodimer, and RAD51 were considered as three individual entities or domains responsible for formation of each FIGNL1 complexes for assessment of binding energetics. All the molecular interactions were analysed and visualized with PyMol 2.2.3 visualization software (https://legacy.ccp4.ac.uk/).

### Protein Purification

Human MMS22L-TONSL (*22*), FIGNL1-dNLS, miBRCA2 and RPA(*69*) were expressed in Spodoptera frugiperda 9 (*Sf*9) insect cells in SFX Insect serum-free medium (Hyclone) using the Bac-to-Bac expression system (Invitrogen), according to manufacturer’s recommendations. MMS22L and TONSL proteins were co-expressed to prepare MMS22L-TONSL protein complex using pFB-MBP-MMS22L-His and pFB-GST-TONSL plasmids. The protein was extracted from *Sf9* cell pellets with 325mM NaCl as described previously(*22*). The protein was then bound to amylose resin (New England Biolabs) via the MBP tag on the MMS22L subunit. The amylose resin with bound protein was washed with Amylose Wash Buffer (50mM Tris– HCl [pH 7.5], 2mM β-mercaptoethanol, 200mM NaCl, 10% glycerol, and 1mM phenylmethanesulfonyl fluoride [PMSF]) and eluted using Amylose Wash Buffer supplemented with 10mM Maltose. Both MBP and GST tags were then cleaved by PreScission protease. The GST-tag was not used during the purification. The final protein complex was obtained by affinity purification on NiNTA agarose (Qiagen) via a 10x-His-tag on the C-terminus of MMS22L. The NiNTA resin bound protein was washed with NiNTA Wash Buffer (50mM Tris–HCl [pH 7.5], 2mM β-mercaptoethanol, 150mM NaCl, 10% glycerol, 1mM PMSF and 40mM imidazole) and eluted using NiNTA Wash Buffer supplemented with 400 mM imidazole. FIGNL1-dNLS was purified from pFB-MBP-FIGNL1-dNLS-His plasmid using identical procedure as MMS22L-TONSL. The miBRCA2 proteins were purified from pFB-2xMBP-Mini_BRCA2-FLAG-His plasmids using similar protocols with 1 M NaCl during the MBP washing steps. The NiNTA resin with bound miBRCA2 was washed with the NiNTA Wash Buffer containing 58mM imidazole and eluted using NiNTA Wash Buffer supplemented with 300mM imidazole. The final protein sample was dialyzed against storage buffer (50mM Tris–HCl [pH 7.5], 5mM β-mercaptoethanol, 150mM NaCl, 10% glycerol, and 0.5mM PMSF) (amino acid sequence of miBRCA2 is provided in the table below).

Human RPA1, RPA2, and RPA3 were co-expressed to produce RPA heterotrimer using pFB-RPA1, pFB-RPA2 and pFB-6xHis-RPA3 was purified by exploiting the His-tag on RPA3 (*69*) using NiNTA affinity chromatography, followed by HiTrap Blue HP, HiTrap desalting and HiTrap Q HP chromatography columns (all GE Healthcare) using ÄKTA pure (GE Healthcare). Wild type RAD51, as well as the RAD51-K133R variant, were expressed from from pMALT-P-RAD51 and pMALT-P-RAD51-K133R plasmids in BL21(DE3)pLysS *E. coli* cells and purified using amylose affinity chromatography (New England Biolabs) followed by Hitrap Q HP chromatography (Cytiva) (*22*). The MBP tag was cleaved during the purification of RAD51 variants. Human DNA2 was expressed from the pFB-His-DNA2-FLAG plasmid in *Sf*9 insect cells and purified by affinity chromatography taking advantage of the N-terminal 6x His-tag and the C-terminal FLAG-tag (*22*).

### DNA Substrate Preparation

For DNA strand exchange assays (*22*), the 3’-tailed DNA was prepared by annealing oligonucleotides RJ-167-mer and RJ-PHIX-42-1 in 1:1 ratio with the annealing buffer [100mM Tris-HCl (pH 8.0), 500mM NaCl and 100mM MgCl_2_). The dsDNA was generated by radio-labeling RJ-Oligo1 with ^32^P at the 3’-end and annealing it to RJ-Oligo2 in 1:2 ratio. DNA protection assays were performed using PC1253 as ssDNA oligonucleotide. DNA binding assays were performed using PC1253 ssDNA oligonucleotide annealed with PC1253c complementary oligonucleotide to form the dsDNA. All oligonucleotides were ^32^P-labeled at the 3ꞌ terminus with [α-^32^P] dCTP (Hartmann Analytic) and terminal transferase (New England Biolabs) according to the manufacturer’s instructions. Sequences of all the oligo substrates used in the study are listed in the table below.

### DNA binding Assay

Binding reactions (15µl volume) were carried out in binding buffer [25mM Tris-acetate (pH 7.5), 3mM EDTA, 1mM dithiothreitol (DTT), and 100 µg/ml bovine serum albumin (BSA)], and DNA substrate (1nM, in molecules). 3mM EDTA was replaced with 1mM ATP (Sigma, A7699) and 2mM magnesium acetate, where indicated. Proteins were added and incubated for 15 minutes on ice. Loading dye (5µl; 50% glycerol [w/vol] and bromophenol blue) was added to the reactions and the products were separated on 6% polyacrylamide gels (ratio acrylamide:bisacrylamide 19:1, Bio-Rad) in TAE (40mM Tris, 20mM acetic acid and 1mM EDTA) buffer at 4 °C. The gels were dried on 17 CHR paper (Whatman), exposed to a storage phosphor screen (GE Healthcare) and scanned by a Typhoon Phosphor Imager (FLA9500, GE Healthcare). Signals were quantified using ImageJ and plotted with GraphPad Prism.

### DNA Strand Exchange Assay

2nM (in molecules) cold 3’ tailed DNA substrate was incubated with the indicated concentration of RAD51 for 5 minutes in a buffer [25mM Tris Acetate (pH 7.5), 1mM DTT, 1mM MgCl_2_, 2mM ATP, 2mM CaCl_2_, and 0.1 mg/ml Recombinant Albumin (NEB) in a total reaction volume of 15 µl. FIGNL1-dNLS was titrated on top and incubated at 37°C for another 5 minutes. This was followed by adding 2nM 32P-radiolabeled dsDNA (in molecules) and incubation for 30 minutes at 37°C. The reaction was stopped with 5 µl of stop solution (150mM EDTA, 0.2% sodium dodecyl sulfate, 30% glycerol and bromophenol blue) and 1µl Proteinase K (Roche) for another 10 minutes at 37°C. Reaction products were separated by running 6% PAGE gel in 1X TAE buffer for 70 minutes at 60 V. For reactions with RPA, 50nM RPA was preincubated with the cold substrate for 5 minutes at 37°C before adding RAD51. For reactions with miBRCA2, 30nM of miBRCA2 was incubated with RAD51 for 5 minutes at 37°C. Where indicated, MMS22L-TONSL was incubated after adding RAD51 for 5 minutes at 37°C before adding FIGNL1-dNLS. Where proteins were omitted, protein dilution buffers were used instead [25mM Tris-Cl (pH 7.5) and 100mM NaCl]. The gels were dried, exposed to a Phosphor imager screen, and analyzed by Typhoon FLA 9500.

### DNA protection assays

DNA2-catalyzed nuclease assays were performed in a 15μl volume reaction buffer [25mM Tris-acetate (pH 7.5), 2mM magnesium acetate, 1mM ATP, 1mM DTT, 0.1 mg/ml BSA, 1mM phosphoenolpyruvate (PEP), and 80U/ml pyruvate kinase (Sigma)], and 1nM substrate (in molecules). RAD51-WT (as indicated in the figure) was preincubated with the substrates for 15 minutes at 37°C in the reaction buffer, which was then supplemented with indicated amounts of DNA2 and FIGNL1-dNLS and the reaction was continued for 30 minutes at 37°C. Reactions were stopped by adding 0.5μl of 0.5M EDTA and 1μl Proteinase K (Roche, 18 mg/ml), and incubated at 50 °C for 30min. An equal amount of formamide dye (95% [v/v] formamide, 20mM EDTA, bromophenol blue) was added, samples were heated at 95°C for 4min and separated on 15% denaturing polyacrylamide gels (ratio acrylamide:bis acrylamide 19:1 [Biorad]). After fixing in a solution (40% methanol, 10% acetic acid and 5% glycerol) for 30 min, the gels were dried on 3MM paper (Whatman), exposed to storage phosphor screens (GE Healthcare) and scanned with a Typhoon 9500 Phosphor Imager (GE Healthcare).

### DR-GFP Assay

U2-OS cells with stable integration of DR-GFP were transfected with respective siRNAs for 24 hours followed by ISceI expressing pCBASceI transfection for further 48 hours. Cells were then harvested, washed with PBS+10% FBS and filtered through 40µm cell strainer to prepare single cells suspension. Samples were then measured for GFP in BD LSR Fortessa and analyzed by FlowJo software v10.5.0. Remaining samples were then lysed and analyzed by western blotting for respective protein knockdowns.

### Genome Stability Assay

The genome stability assay was carried out according to the standard protocol described previously (*70*). Cells were treated with 500nM Olaparib for 24 hours or 750nM cisplatin for 6 hours. Cells that were treated with Olaparib were exposed to colcemid containing media 12 hours after the start of the treatment and continued the Olaparib and colcemid treatment for the next 12 hours. For the cells that were treated with cisplatin, the drug treated media was washed off and the cells were exposed to colcemid containing media for the next 12 hours. Metaphase spreads were prepared by conventional methods. Metaphase slides were washed with 2x SSC, PBS and MgCl_2_ in PBS, denatured at 80°C after which telomere probes were hybridized onto the metaphase spreads. After hybridization, the slides were washed with 50% formamide in 2x SSC, 0,1X SSC and 4x SSC containing 0,1% Tween-20. The metaphase spreads were stained with DAPI and mounted with MOWIOL. A minimum of 50 metaphase images were captured using Metafer5 and analyzed with Adobe Photoshop for chromosomal aberrations.

### Survival Assay

For the mESCs ten thousand cells were seeded per well in a gelatinized 96-well plate. After 24 h, cells were treated with varying concentrations of olaparib or cisplatin for 72 h. After 72 h, cells were rinsed with PBS and incubated with 1mg/ml of XTT (2,3-bis(2-methoxy-4-nitro-5-sulfophenyl)-5-[(phenylamino)carbonyl]-2H-tetrazolium hydroxide), 2mM phenazine methosulfate (PMS) at 37°C for 60 min. Plates were read in iMark Microplate reader (Bio-Rad) at 455nm.

1×10^3^ mouse mammary tumor cells were seeded in a 6-well plate and 24 hours later cells were treated with indicated concentrations or Olaparib or cisplatin. Cells treated with Olaparib were refreshed with fresh Olaparib every 48 hours for 7 days, while cells treated with cisplatin were refreshed after 12 hours and incubated in fresh media without cisplatin for 7 days. After 7 days, survivability was quantified with CellTiter-Blue® Cell Viability Assay (Promega) as per manufacturer’s instructions and plates were fixed and stained with Brilliant Blue G (Sigma Aldrich).

### Interaction assays

To study the interaction between FIGNL1-dNLS and MMS22L-TONSL in the presence of RAD51-K133R mutant, soluble extracts of GST-FIGNL1-dNLS prepared from pGEX-2T-FIGNL1-dNLS plasmid in Rosetta (DE3)pLysS *E. Coli* cells. Each culture was supplemented with 0.2% glucose, induced with 0.5 mM IPTG and grown overnight at 18 °C. The cells were then pelleted at 2,500 g for 15 minutes at 4°C, washed once with STE buffer (10mM Tris-HCl pH 8, 500mM NaCl, 1 mM EDTA), snap-frozen and kept at −80 C until use. The soluble extract containing GST-FIGNL1-dNLS was prepared by resuspending the bacterial pellets in GST-buffer (50mM Tris HCl pH 7.5, 5mM MgCl_2_, 2mM ATP, 1mM PMSF, 2mM β-ME, 150mM NaCl 10% Glycerol, 1mM EDTA, 1 mM) followed by sonication and centrifugation for 30min at 48000g. The supernatant was immobilized on GST resin (GE Healthcare) upon incubation for 1 hour at 4 °C with continuous rotation. The resin was washed 3 times with GST-buffer post incubation. Following this, either 50µl binding buffer (50mM Tris–HCl, pH 7.5, 2mM β-mercaptoethanol, 3mM EDTA, 100mM NaCl, 0.2µg/µl bovine serum albumin [BSA], 1mM PMSF) alone or that supplemented with 0.5µg of RAD51-K133R or RAD51-K133R (pre-bound with ssDNA [PC1253]) was added to the immobilized GST-FIGNL1-dNLS for 1 hour at 4 °C with continuous rotation. ssDNA was pre-bound for 1 hour to RAD51-K133R at 37 °C. Following this, the resin was washed once with GST buffer and MMS22L-TONSL was added to the appropriate samples and incubated again for 1 hour, followed by three washes and elution using 10mM reduced glutathione (Sigma). The eluates were separated on a 10% SDS-PAGE gel and proteins were detected by western blotting using indicated primary antibodies (table below).

### GFPtrap pull down

20×10^6^ EGFP-FIGNL1 over expressing RPE1 cells were harvested and subjected to nuclear fractionation. Briefly, cells were lysed with cytoplasmic extraction buffer (10mM HEPES, 1.5mM MgCl_2_, 10mM KCl, 0.5mM DTT, 0.05% NP40 supplemented with protease inhibitor cocktail (PIC)) and incubated ice for 10 minutes. Cytoplasmic fraction was then separated by centrifugation at 3,000rpm for 5 minutes. Nuclear fraction at the pellet was then separated and lysed with immunoprecipitation buffer (50mM Tris-HCl (pH-7.4), 150mM NaCl, 10% glycerol, 1% Triton X-100, 0.5mM EDTA, 0.5mM EGTA, and Protease Inhibitor cocktail) along with Dounce homogenization with 25 strokes of up and down, on ice. Finally, the lysed nuclear fraction was collected by centrifugation at 13,000rpm for 10 minutes. Cleared nuclear fraction was quantified by BCA method (Pierce, Invitrogen) and used for overnight GFPtrap magnetic agarose bead pull-down (Proteintech) after separating 5% lysate as input. After overnight GFPtrap pull down, beads were washed 5 times with wash buffer (10 mM Tris-HCl pH 7.5, 150 mM NaCl, 0.05 % NP40 Substitute, 0.5 mM EDTA supplemented with PIC). Pull down eluates were separated by heating with NuPAGE^TM^ LDS sample buffer (Invitrogen) + 4x DTT at 95°C for 15 minutes and analyzed by western blotting.

**Table:**
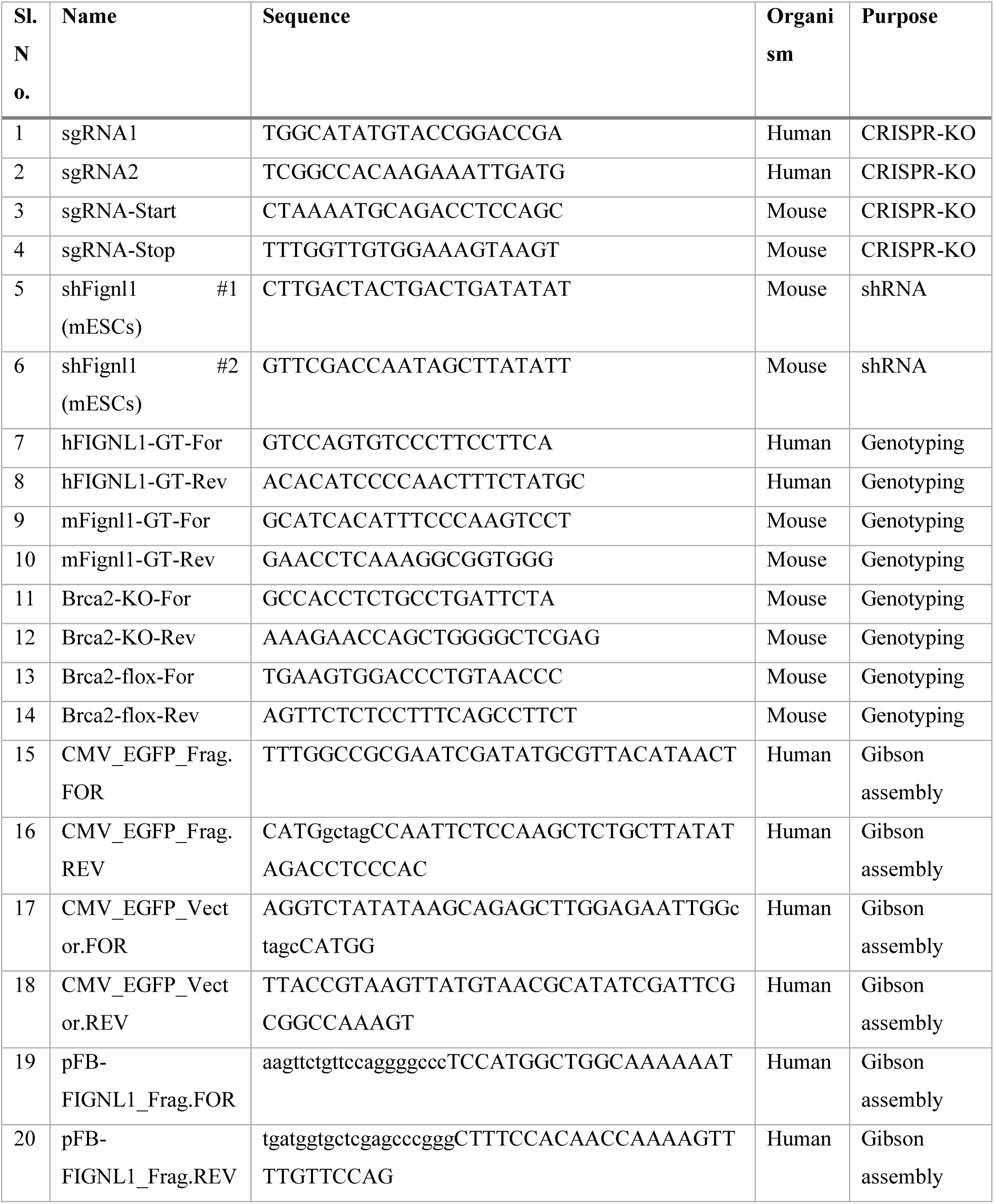
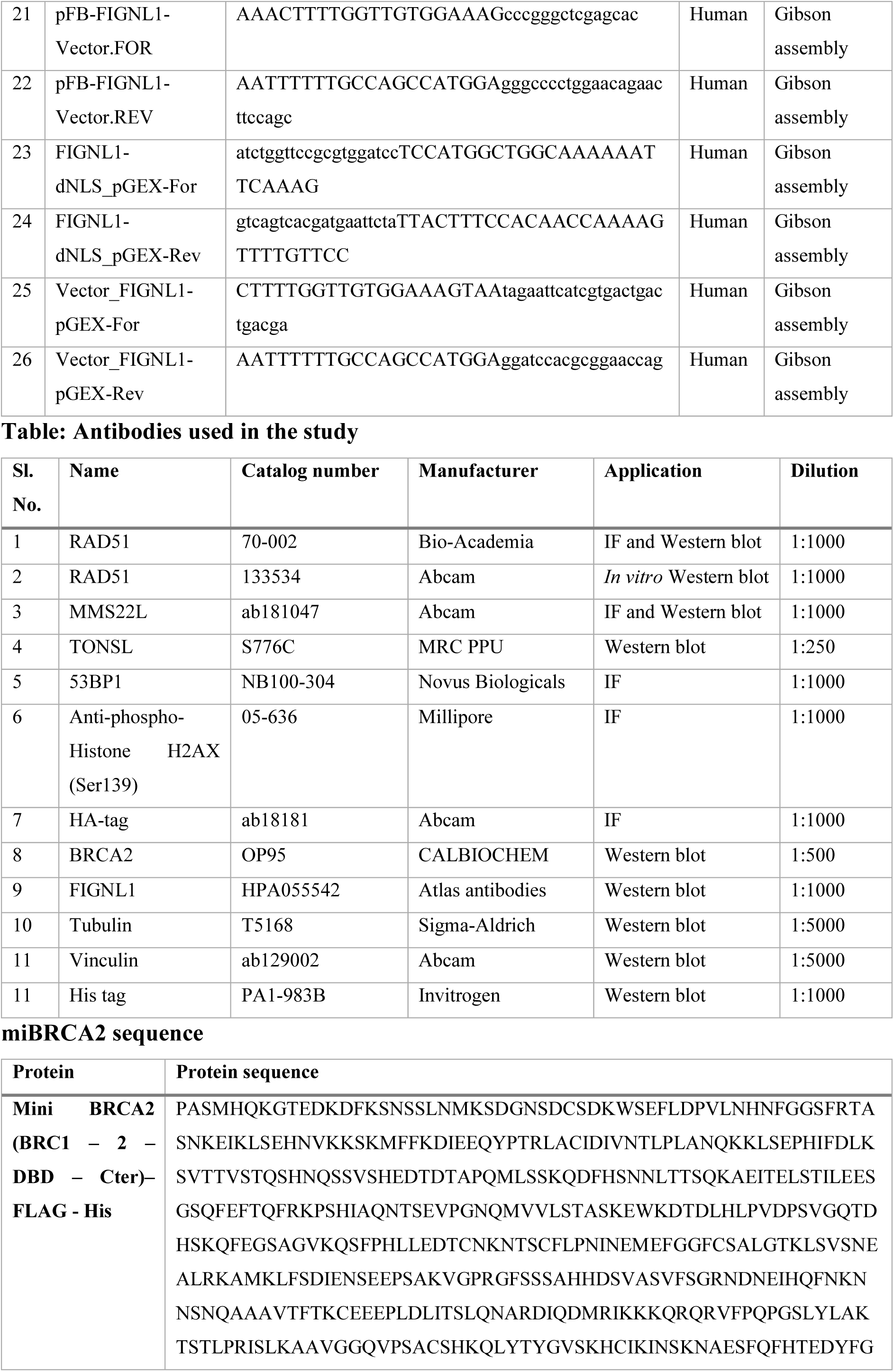
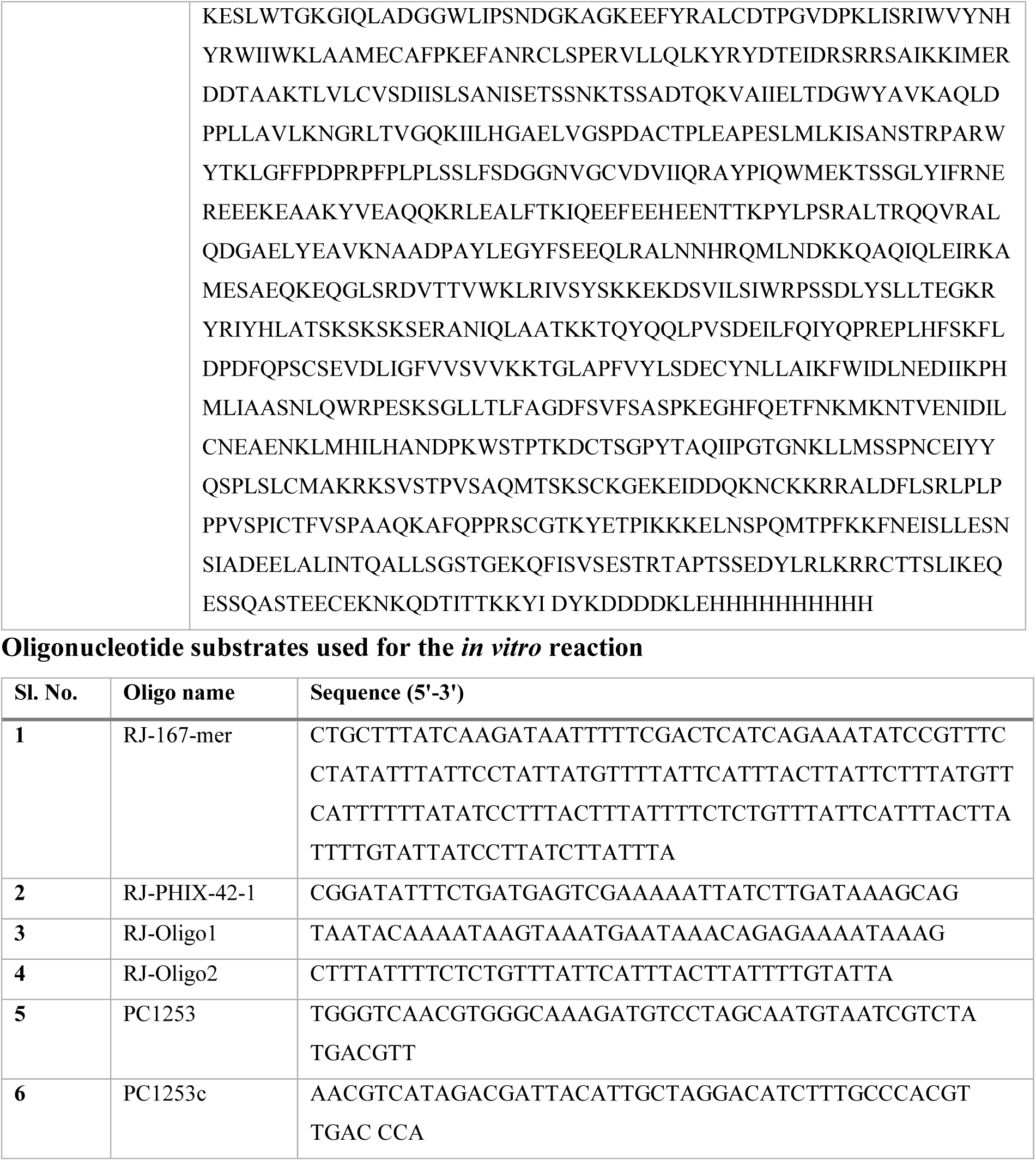
Oligos used in the study.

---

**Supplementary Figure 1:**
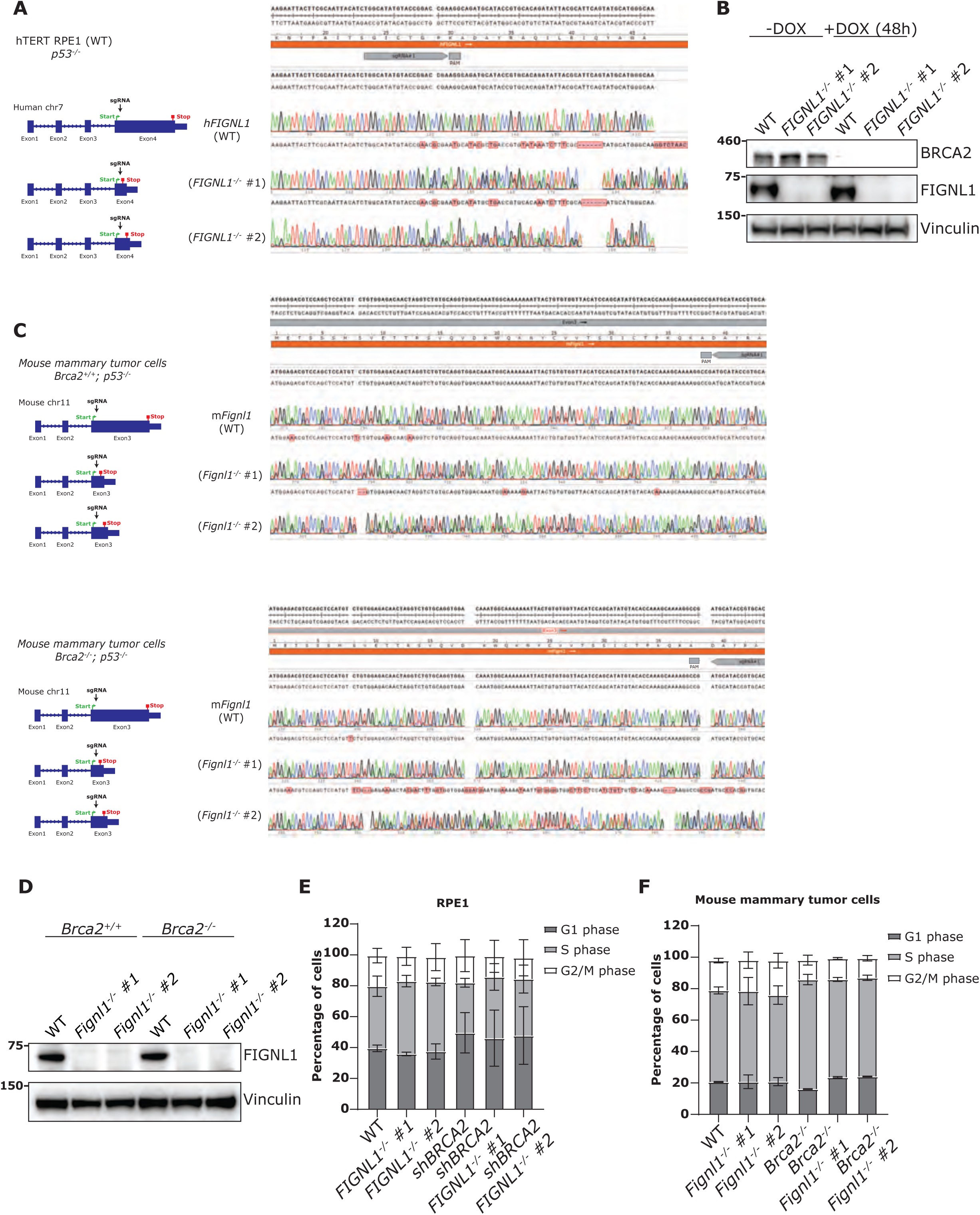
Validation of FIGNL1 knock-out cells in human RPE1 and mouse mammary tumor cells. (A) Left: Schematics of the results of knocking-out *FIGNL1* in RPE1-WT cells. Right: Genotyping results from RPE1-WT and 2 different knockout clones of *FIGNL1*. Electropherogram representative figure showing the alignment of the *FIGNL1* genomic DNA region at the indicated binding site of sgRNA followed by its PAM sequence. (B) Western blot analysis of BRCA2 with doxycycline induced knock-down and FIGNL1 levels upon CRISPR-Cas9 mediated knockout in RPE1 cells. (C) Left: Schematics of the results of knocking-out *Fignl1* in *Brca2* proficient (top) and deficient (bottom) mouse mammary tumor cells. Right: Genotyping results from *Brca2* proficient (top) and deficient (bottom) mouse mammary tumor cells with 2 different clones of *Fignl1* knockouts each. Electropherogram representative figure showing the alignment of *Fignl1* genomic DNA region at the indicated binding site of sgRNA followed by its PAM sequence. (D) Western blot analysis of FIGNL1 levels from each clone upon CRISPR-Cas9 mediated knockout in mouse mammary tumor cells. (E) The percentage of cells in the different phases of the cell cycle profile of RPE1 and RPE1 *FIGNL1*-KO cells. Error bars are indicative of mean±SD from 2 independent flow cytometry experiments. (F) The percentage of cells in the different phases of the cell cycle profile of mouse mammary tumor cells with *Fignl1*-KO. Error bars are indicative of mean±SD from 2 independent flow cytometry experiments.

**Supplementary Figure 2:**
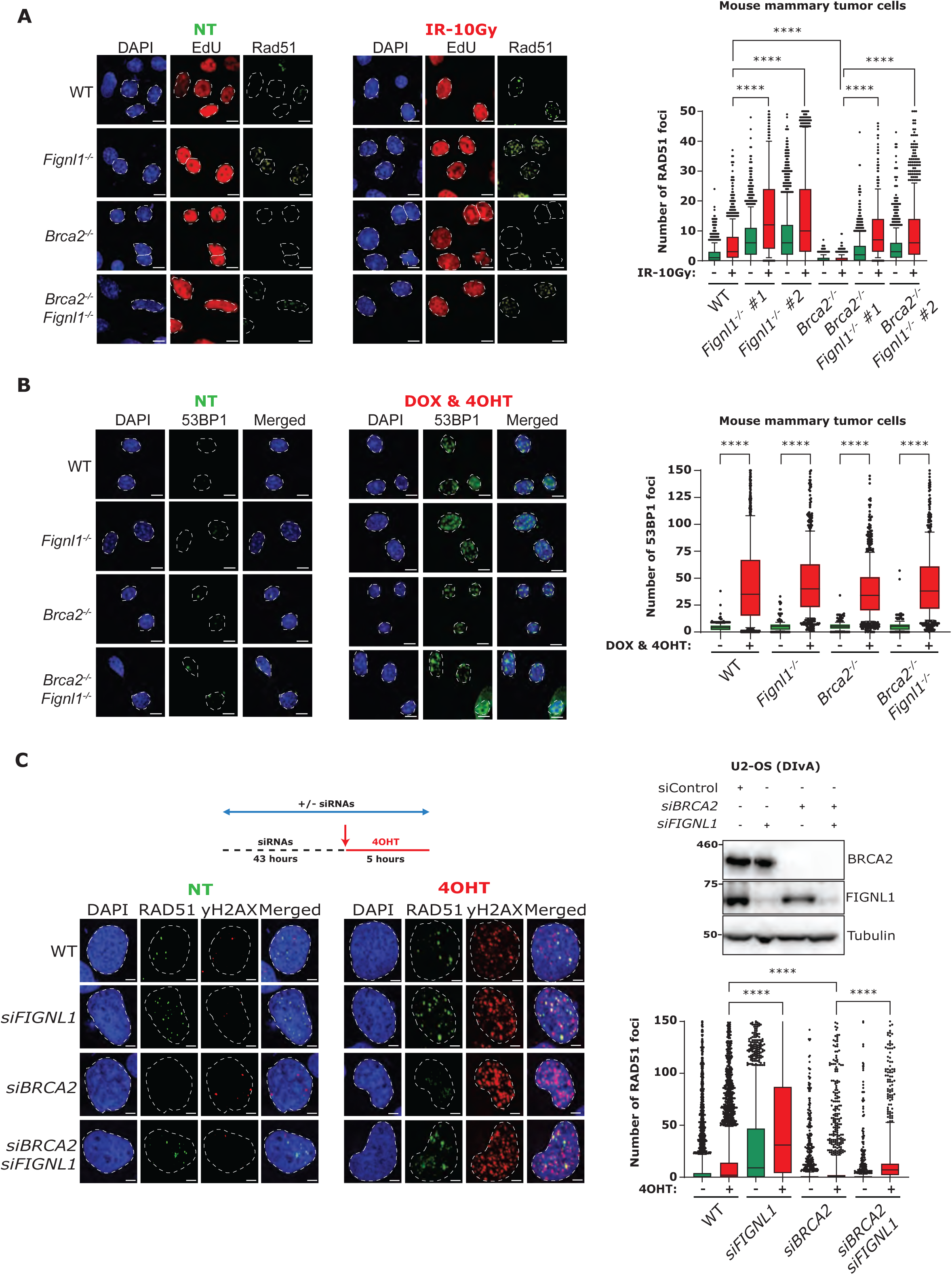
Validation of RAD51 rescue in BRCA2-FIGNL1 deficient cells in mouse mammary tumor cells and human U2-OS cells. (A) Left: representative images of high-content, automated RAD51-IF images showing the EdU and RAD51 foci in untreated and ionizing radiation treated mouse mammary tumor cells; scale bar: 20µm. Right: Automated QIBC plots of the RAD51 foci per nucleus; scatter box plot showing the foci distribution between 10-90th percentile from 1500 cells (experiment was repeated 3 times). P-values from Mann-Whitney test (two-tailed) comparing between indicated groups (****: <0.0001). (B) Left: representative images of 53BP1-IF images showing the 53BP1 foci in untreated and AsiSI induced mouse mammary tumor cells; scale bar: 20µm. Right: Automated QIBC plots of the 53BP1 foci per nucleus; scatter box plot showing the foci distribution between 10-90th percentile from 1500 cells (experiment was repeated 3 times). P-values from Mann-Whitney test (two-tailed) comparing between indicated groups (****: <0.0001). (C) Top Left: Schematics showing the experimental setup for RAD51-IF in U2-OS AsiSI (DIvA) cells. 48 hours of siRNA transfection to knock-down BRCA2 and FIGNL1 with last 5 hours to translocate constitutively expressing cytoplasmic AsiSI to the nucleus by addition of 4-hydroxytamoxifen (4-OHT). Top right: Western blot analysis showing the levels of BRCA2 and FIGNL1 upon siRNA mediated knock-down in DIvA cells. Bottom left: representative images of high-content, automated RAD51-IF images showing the RAD51 and γH2AX foci in untreated and AsiSI induced DIvA cells; scale bar: 5μm. Bottom right: Automated QIBC plots of the RAD51 foci per nucleus; scatter box plot showing the foci distribution between 10-90th percentile from 1500 cells (experiment was repeated 3 times). P-values from Mann-Whitney test (two-tailed) comparing between indicated groups (****: <0.0001).

**Supplementary Figure 3:**
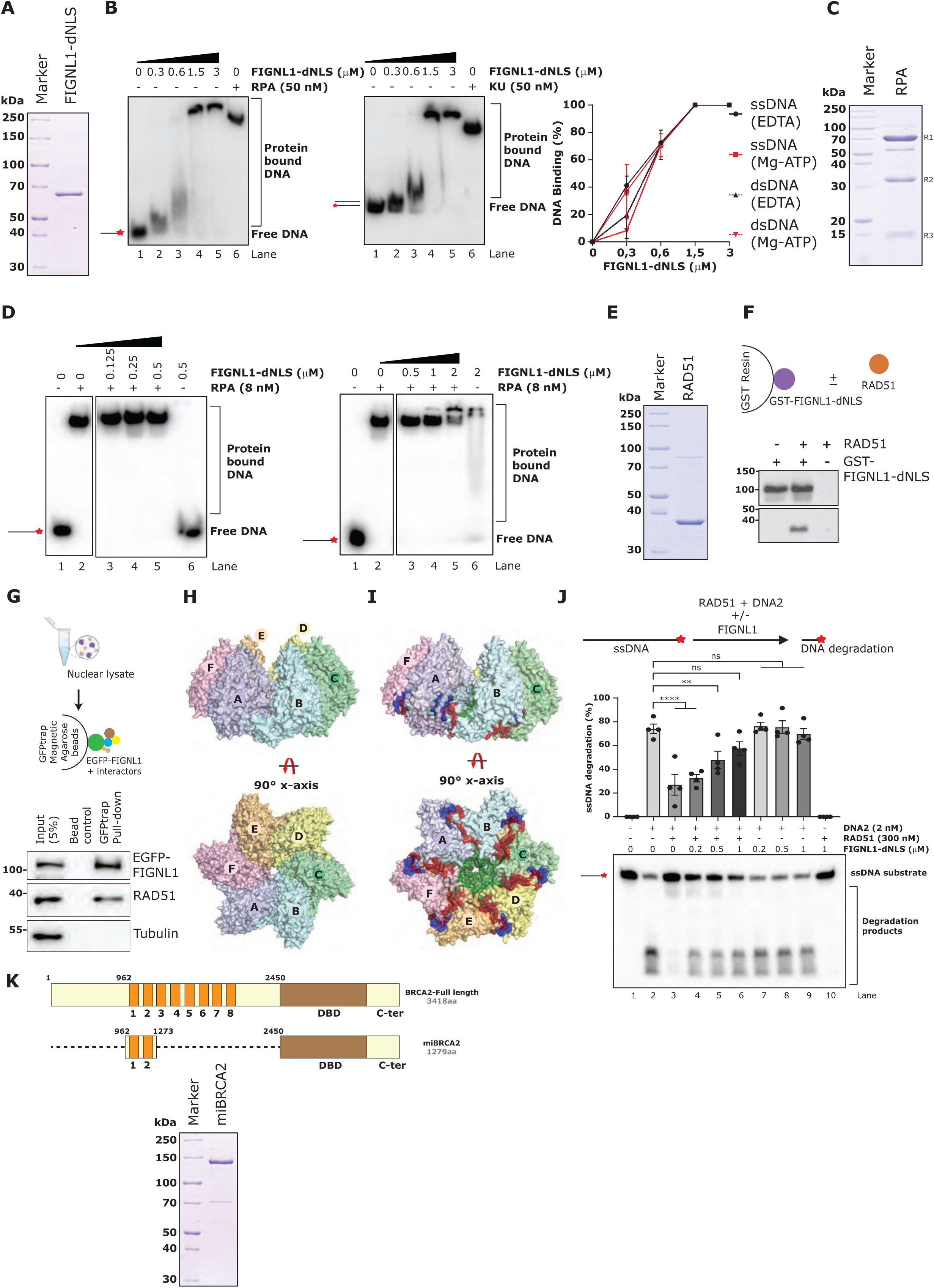
FIGNL1 dissociates RAD51 from ssDNA. (A) Recombinant FIGNL1-dNLS protein purified from baculovirus infected insect *Sf9* cells. The purified protein was separated on a polyacrylamide gel and stained with Coomassie blue. (B) Representative images of electrophoretic mobility shift assay showing the concentration dependent binding to ssDNA (left) or dsDNA (middle); RPA was used as a control for ssDNA binding and yeast Ku as a control for dsDNA binding. The red asterisk indicates the position of the radioactive label. Right: quantification of binding efficiency with ssDNA and dsDNA in the presence or absence of Mg2+ and ATP; error bars are indicative of mean±SEM. (C) Recombinant RPA protein purified from baculovirus infected insect *Sf9* cells. The purified protein was separated on a polyacrylamide gel and stained with Coomassie blue. R1: RPA1, R2: RPA2 and R3: RPA3. (D) Representative images of electrophoretic mobility shift assay showing the concentration dependent DNA binding efficiency of FIGNL1-dNLS in the presence of 8nM RPA; Left: low concentration, Right: high concentration of FIGNL1-dNLS. The red asterisk indicates the position of the radioactive label. (E) Recombinant RAD51 protein purified from BL21-(DE3)pLysS *E. coli* cells. The purified protein was separated on a polyacrylamide gel and stained with Coomassie blue. (F) Top: Schematics of the in vitro interaction assay between FIGNL1-dNLS (immobilized) and RAD51. Bottom: Western blot results indicative of the interaction between FIGNL1 and RAD51 in the *in vitro* interaction assay. (G) Top: Schematics of the GFPtrap pull-down of the EGFP-FIGNL1 and RAD51 from nuclear extracts. EGFP-FIGNL1 stably over-expressing cells in *FIGNL1^-/-^* RPE1 cells were used for nuclear fractionation and GFPtrap pulled down. Bottom: Western blot results indicative of the interaction between FIGNL1 and RAD51. (H) The side and top view of the FIGNL1 hexamer. The monomers are shown as surface representation (Chain A: Light blue; Chain B: Pale cyan; Chain C: Pale green; Chain D: Light yellow; Chain E: Light Orange; Chain F: Light pink). (I) The top and side view of FIGNL1 showing the interacting domains participating in formation of the hexamer (N-terminal: Blue, ATPase: Dark green, and C-terminal: maroon). (J) Top: Quantification of ssDNA degradation was done by normalizing the ssDNA substrate band intensity against the band intensity of the no-protein product; error bars are indicative of mean±SEM from 4 independent experiments (n=4). P-values from Ordinary One-Way ANOVA comparing between indicated groups (**: <0.01; ****: <0.0001). The red asterisk indicates the position of the radioactive label. Bottom: representative gel picture for ssDNA protection assay showing the protection efficiency of RAD51 against degradation by DNA2 nuclease in the presence of increasing concentrations of FIGNL1-dNLS. (K) Top: Schematics of miBRCA2 constructed with BRC repeats BRC1 & 2, DNA binding domain (DBD) and C-terminal domain (C-ter) compared to the full-length BRCA2. Recombinant miBRCA2 protein purified from baculovirus infected insect *Sf9* cells. The purified protein was separated on a polyacrylamide gel and stained with Coomassie blue.

**Supplementary Figure 4:**
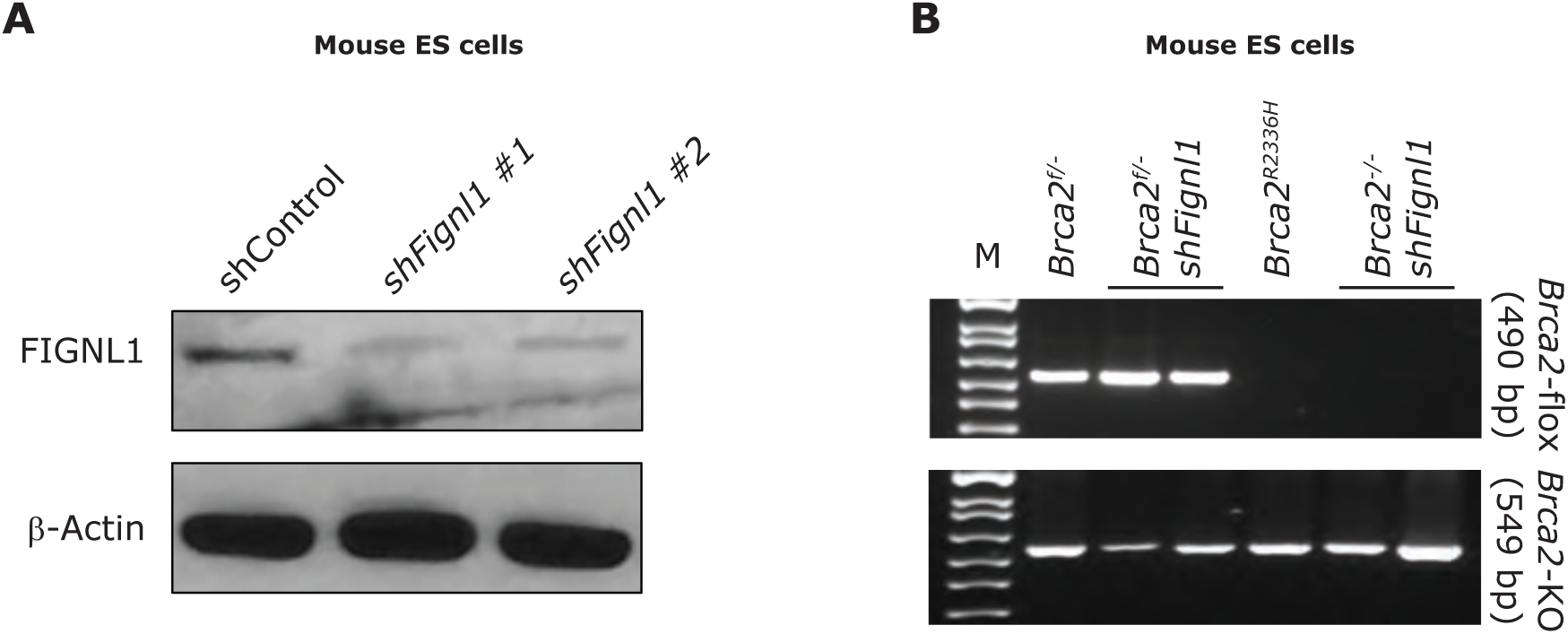
Validation of FIGNL1 knock-down in mouse ES cells. (A) Representative western blot images showing the FIGNL1 knock-down in mouse embryonic stem cells. (B) Representative PCR genotyping images of mouse embryonic stem cells with conditional knock-out of BRCA2. Upper lane for floxed allele of BRCA2 (490 bp PCR product) and lower lane for the deleted allele of BRCA2 (549 bp PCR product).

**Supplementary Figure 5:**
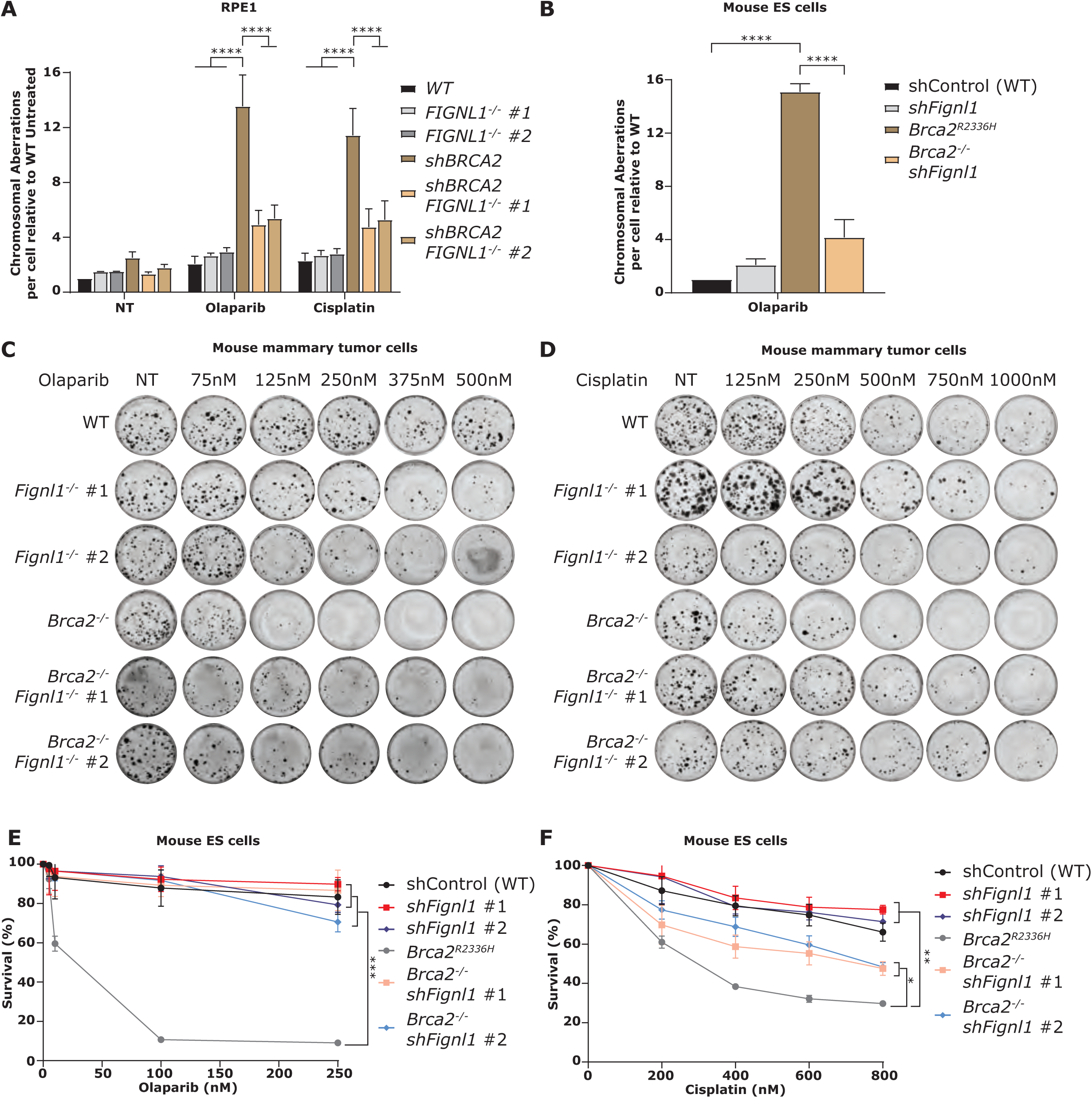
Loss of FIGNL1 in BRCA2 deficient cells reverses genome instability and confers chemoresistance. (A) Quantification of chromosomal aberrations in *FIGNL1^-/-^* RPE1 clones in the presence or absence of BRCA2 in response to Olaparib and cisplatin from 3 independent experiments in which 50 individual metaphases were analyzed for each sample per experiment. P values from Ordinary two-way ANOVA (****: <0.0001) (B) Quantification of chromosomal aberrations in mouse embryonic stem cells with FIGNL1 knock down clones in the presence or absence of *Brca2* in response to Olaparib from 3 independent experiments in which 50 individual metaphases were analyzed for each sample per experiment. P values from Ordinary two-way ANOVA (****: <0.0001) (C) Representative clonogenic images of mouse mammary tumor cells in response to Olaparib. (D) Representative clonogenic images of mouse mammary tumor cells in response to cisplatin. (E) Percentage survivability of mouse embryonic stem cells in response to Olaparib; quantification was performed by normalization to non-treated cells. P-values from Dunnett’s multiple comparison One-way ANOVA comparing between indicated groups (*: <0.05, ***: <0.001). (F) Percentage survivability of mouse embryonic stem cells in response to cisplatin; quantification was performed by normalization to non-treated cells. P-values from Dunnett’s multiple comparison One-way ANOVA comparing between indicated groups (*: <0.05, **: <0.01).

**Supplementary Figure 6:**
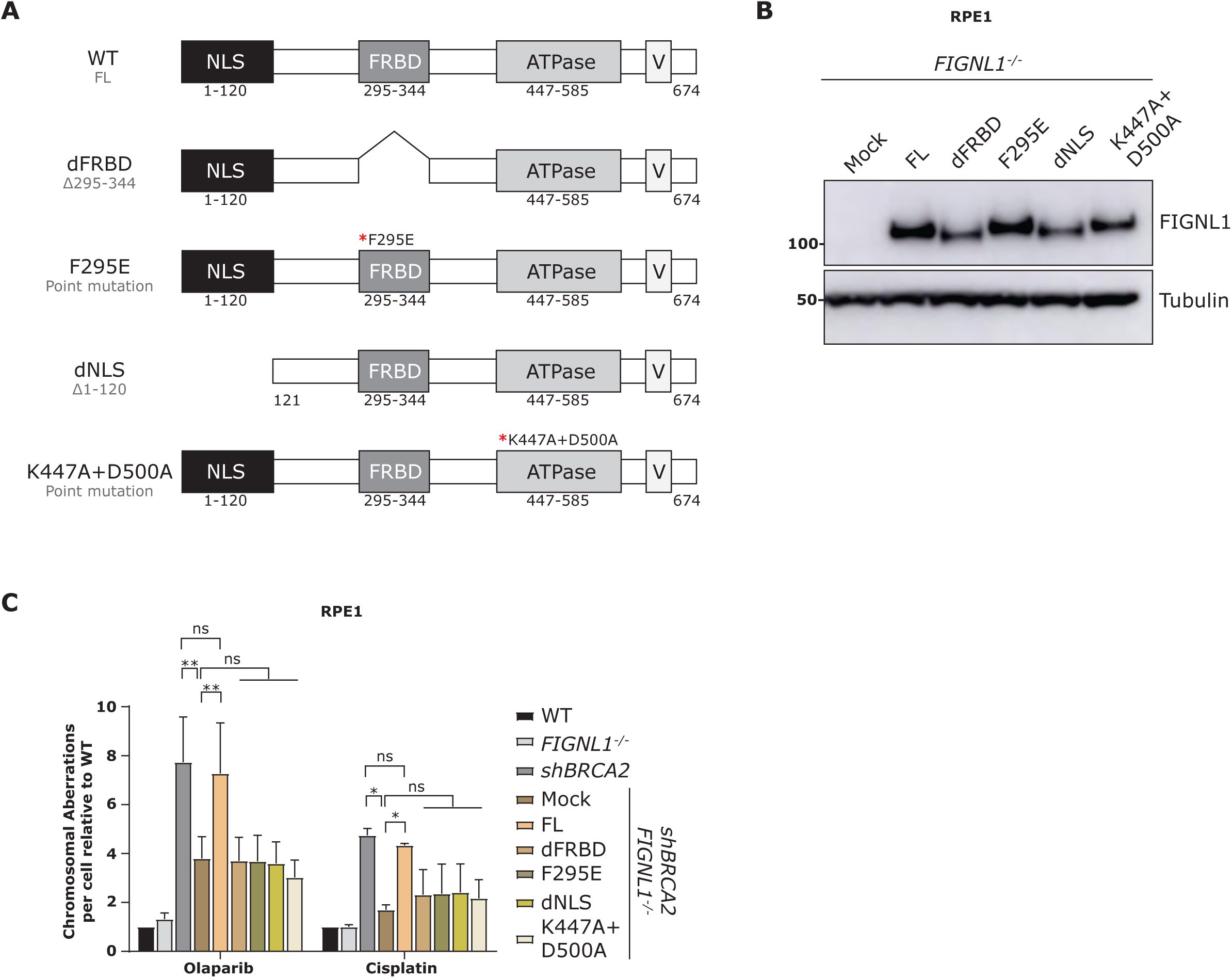
FIGNL1-RAD51 interaction is essential for genome stability. (A) Schematics of the FIGNL1 mutant proteins (B) Western blot analysis of FIGNL1 mutants fusion with GFP protein. (C) Quantification of chromosomal aberrations in *FIGNL1^-/-^* RPE1 cells reconstituted with FIGNL1 mutants in response to Olaparib and cisplatin from 3 independent experiments in which 50 individual metaphases were analyzed for each sample per experiment. P values from Ordinary two-way ANOVA (ns-nonsignificant; *: <0.05; **: <0.01).

**Supplementary Figure 7:**
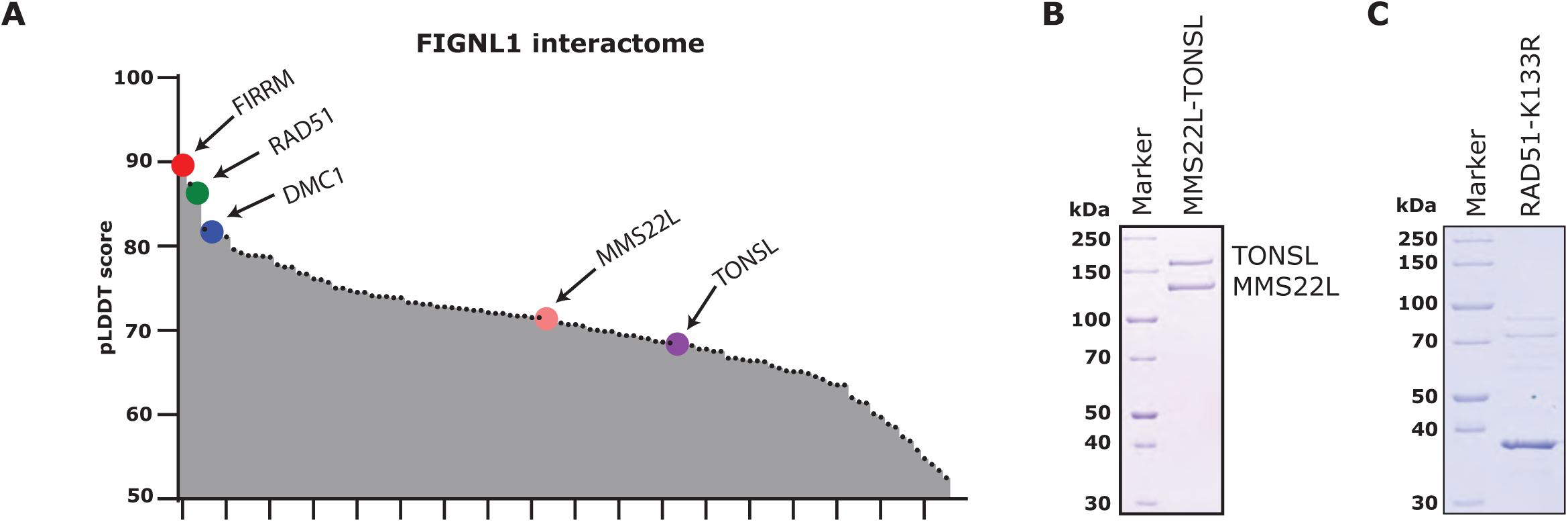
Identification of MMS22L-TONSL complex as FIGNL1 interacting partners. (A) Predictome analysis highlighting the FIGNL1 interacting partners involved in genome maintenance. (B) Recombinant MMS22L and TONSL heterodimer purified from baculovirus infected insect *Sf9* cells. Purified proteins were separated on a polyacrylamide gel and stained with Coomassie blue. (C) Recombinant RAD51-K133R protein purified from BL21-(DE3)pLysS strain in *E. coli* cells. Purified proteins were separated on a polyacrylamide gel and stained with Coomassie blue.

**Supplementary Figure 8:**
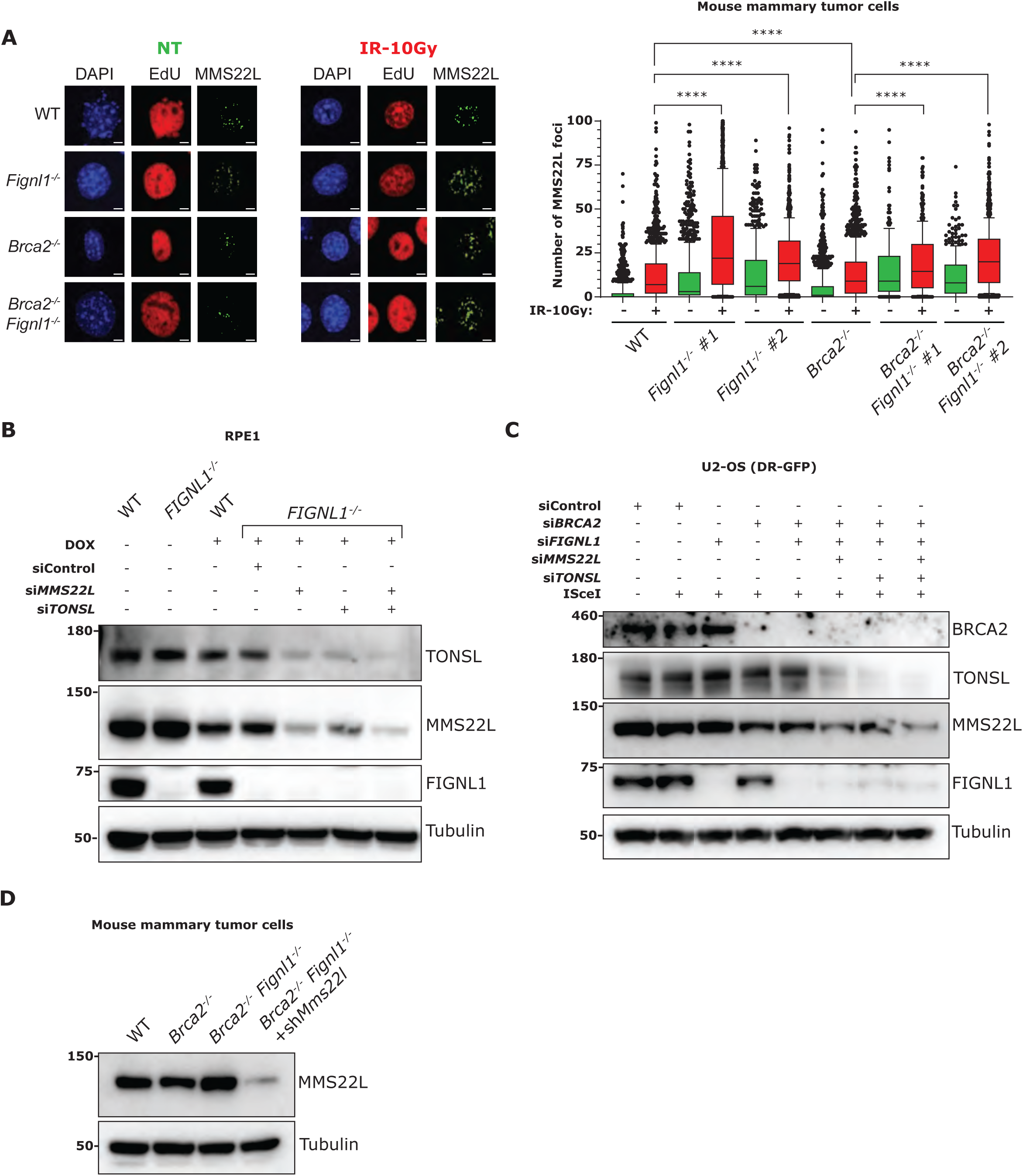
Validation of MMS22L-TONSL levels in the presence or absence of BRCA2 and FIGNL1. (A) Left: Representative high-content, automated IF images showing EdU and MMS22L foci in untreated and ionizing radiation treated cells; scale bar: 5µm. Right: Automated QIBC plots of the MMS22L foci per nucleus; scatter box plot showing the foci distribution between 10-90th percentile from 1500 cells (experiment was repeated 3 times). P-values from Mann-Whitney test (two-tailed) comparing between indicated groups (****: <0.0001). (B) Representative western blot analysis showing the levels of MMS22L, TONSL and FIGNL1 in human RPE1 cells used for RAD51-IF and metaphase spreads. (C) Western blot analysis showing the levels of FIGNL1, BRCA2, MMS22L and TONSL from the corresponding cells used for HR assay. (D) Western blot analysis showing the levels of MMS22L from the corresponding cells used for clonogenic assays in Figure 4F in mouse mammary tumor cells.

